# Predictive Sequence Learning in the Hippocampal Formation

**DOI:** 10.1101/2022.05.19.492731

**Authors:** Yusi Chen, Huanqiu Zhang, Mia Cameron, Terrrence Sejnowski

## Abstract

The hippocampus receives sequences of sensory inputs from the cortex during exploration and encodes the sequences with millisecond precision. We developed a predictive autoencoder model of the hippocampus including the trisynaptic and monosynaptic circuits from the entorhinal cortex (EC). CA3 was trained as a self-supervised recurrent neural network to predict its next input. We confirmed that CA3 is prediction ahead by analyzing the spike coupling between simultaneously recorded neurons in the dentate gyrus, CA3 and CA1 of the mouse hippocampus. In the model, CA1 neurons signal prediction errors by comparing the prediction from CA3 to the next input directly from the EC. The model exhibits the rapid appearance and the slow fading of CA1 place cells, and displays replay and phase precession from CA3. The model could be learnt in a biologically plausible way with the help of error-encoding neurons. Similarities between the circuits in the hippocampus and thalamocortical circuits suggest that such computation motif could also underlie self-supervised sequence learning in the cortex.

## Introduction

The representation of sensory information in cortical structures is encoded in the spatiotemporal patterns of spikes in populations of neurons. During locomotion, the spike timing of neurons in area CA1 of the hippocampus precesses relative to the local phase of the theta wave (O’Keefe & Recce, 1993; Patel et al., 2012). Spike timing is also precisely regulated at the millisecond level to engage spike-timing dependent plasticity (STDP) (Markram et al., 1997; Bi & Poo, 1998). This regulation must take into account time delays for both the conduction of spikes between neurons and transmission delays at synapses. We focus here on the functional implications of this precision for how temporal sequences of spikes are shaped by neural circuits. We show how the temporal precision of spike timing coupled with anatomical wiring could support the learning and replay of temporal sequences in the hippocampal formation.

Cognitive maps are created in the hippocampus, with place cells in rodents responding not only to locomotion signals (O’Keefe & Nadel, 1978) but also to other sensory stimuli, such as reward (Gauthier & Tank, 2018), auditory tones (Aronov et al., 2017), odors and time (Pastalkova et al., 2008; Buzsáki & Tingley, 2018; Eichenbaum et al., 1987). These stimuli are high-dimensional and highly redundant, yet only a few hippocampal neurons are reliably and repetitively activated in a short time interval, forming a relatively low-dimensional dynamical trajectory in activity space (Nieh et al., 2021). The hippocampus therefore learns how to encode high-dimensional sensory and motor signals at the apex of cortical hierarchies into low-dimensional, latent, non-redundant, sequential representations that ultimately support abstract representational learning. After learning sequences of events, the hippocampus then replays them during sleep and immobility when external inputs to the cerebral cortex are suppressed (Wilson & McNaughton, 1994; Skaggs et al., 1992).

Existing computational frameworks (Stachenfeld et al., 2017; Whittington et al., 2020; McNamee et al., 2021; George et al., 2021) have successfully modeled cognitive functions of the hippocampus and reproduced the statistics of place cell under various task conditions. However, these models do not provide an implementation of these cognitive functions based on neural mechanisms or account for the distinct encoding and firing properties of neurons in CA3, CA1, and the dentate gyrus (DG) (Dong et al., 2021; Shin et al., 2022; Lee et al., 2004b,a). For example, CA1 neurons are more responsive to unexpected signals than neurons in other hippocampal areas (Kumaran & Maguire, 2006; Knight, 1996; Duncan et al., 2012) and their activity decays over a time scale of weeks in familiar environments, faster than neuron in other subregions (Fig.S1). In contrast, recurrent circuits in CA3 store an internal representation of sequences, which are regenerating during replay (Wilson & McNaughton, 1994; Skaggs et al., 1992) and preplay (Dragoi & Tonegawa, 2011). Neural place fields emerge faster in CA1 but are generally more stable in CA3 upon remapping (Lee et al., 2004a; Shin et al., 2022). Schapiro et al. (Schapiro et al., 2017) proposed a complementary learning systems for CA1 and CA3 that reconciled statistical learning with episodic memory. We exploit these functional differences for a temporal predictive learning theory of sequences in the hippocampus.

Predictive coding efficiently encodes visual features in lower cortical layers, enabling higher layers to represent more abstract features (Lotter et al., 2020; Rao & Ballard, 1999). This study builds upon these findings by extending predictive coding into the temporal domain to model interactions among hippocampal subregions. We confirmed the temporal prediction hypothesis by analyzing neural recordings. We were unable to replicate experimental findings with a recurrent network model of CA3 that simply learned sequences. However, when we trained a network to make temporal predictions, we were able to successfully replicate the observed statistics of neural activity, the qualitatively distinct dynamics in CA1 and CA3 place cells, and representational learning to generate replay. We further demonstrated a biologically plausible predictive learning rule.

## Results

### Temporal Prediction Hypothesis

Figure 1 summarizes the major connectivity in the hippocampal formation. The entorhinal cortex (EC) is the major cortical input to the hippocampus and is the major recipient of its output. Among hippocampal subregions, recurrently connected CA3 is ideal for storing internal states in the form of attractor dynamics (Hopfield, 1982). Area CA1 receives inputs from two pathways projecting from the EC to CA1: an indirect pathway via DG and CA3 and a direct pathway from the EC. Moreover, the two pathways are delayed to different extents because there are more synaptic delays in the indirect path through CA3 (Sabatini & Regehr, 1999). Assuming a synaptic transmission delay *τ >* 0, signals transmitted through the indirect pathway to CA1 are delayed by 3*τ* while those going through the direct pathway are only delayed by *τ*. An *in vitro* electrophysiology study (Leung et al., 1995) measured a 2.5 ms delay from EC to CA1 through the direct pathway and a 9-17 ms delay through the trisynaptic indirect pathway. The delay from EC to DG was 1.7 ms.

**Figure 1:**
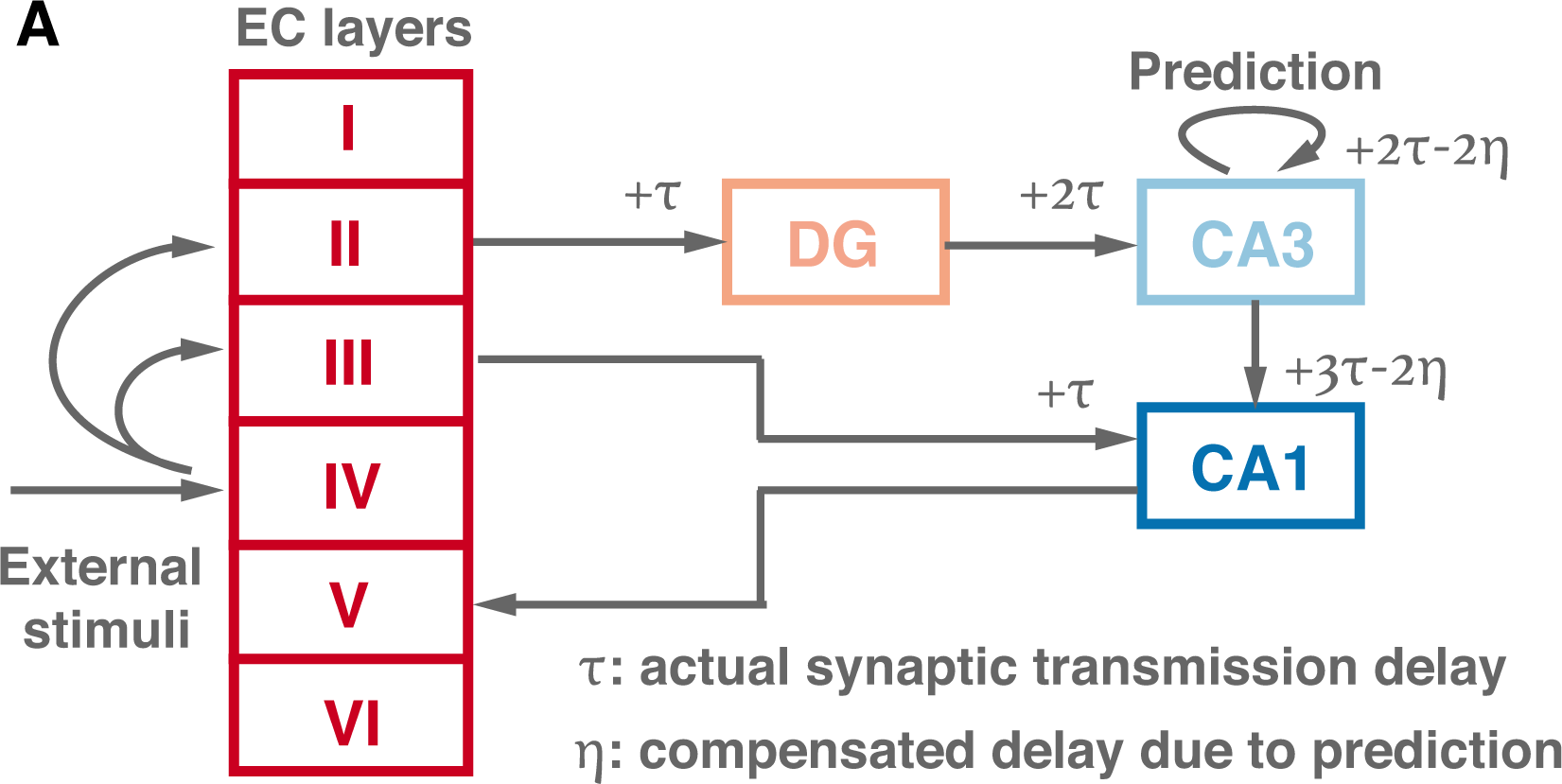
The circuit within the hippocampal formation. (A) Anatomical wiring and interregional delays. External sensory stimuli start from different cortical layers in EC and reach CA1 through two pathways, forming a self-supervised structure. Assume that spikes are delayed by *τ* after one synaptic transmission, they will be delayed by 3*τ* and *τ* at CA1 through the indirect and direction pathway, respectively. We hypothesize that CA3 predicts the future (−2*η*) to compensate for the accumulated transmission time difference (+2*τ*).

The function of this seemingly redundant and asynchronous transmission from EC to CA1 suggests that CA3 may be making predictions about future inputs, which can then be compared at CA1 with the less delayed teacher signal from the direct pathway. This comparison is similar to a Bayesian filter (Wikipedia, 2021) where future predictions based on currently available information are compared with future observations to update the model. Therefore, we hypothesize that prediction errors are computed at CA1 and can be used to refine the internal model stored in CA3. In this way, interactions between the cortex and the hippocampus form a self-supervised loop, which enables the circuit motif to learn and remember the latent variables of a predictive autoencoder represented in CA3 as sequences.

### Neural evidence for transmission delay and predicting ahead

To verify the above hypothesis, we analyzed simultaneously recorded neural activities from these subregions for evidence of transmission delay and predicting ahead. Assuming that neural signal propagation strictly follows the anatomical organization of the hippocampal formation in Fig. 1, signals encoded by a region should be correlated with the upstream signal shifted by an interregional time delay. Ideally, if a location-sensitive neuron in EC has a bell-shaped response curve *f* (*x*), where *x* represents any arbitrary physical variable such as location, its direct downstream DG neuron should exhibit a response *f* (*x − τ*) where *τ* refers to the default interregional delay (Fig. 2A). Similarly, the response curves of their downstream neurons in CA3 and CA1 should be *f* (*x −* 2*τ*) and *f* (*x −* 3*τ*), respectively (dashed lines in Fig. 2A).

**Figure 2:**
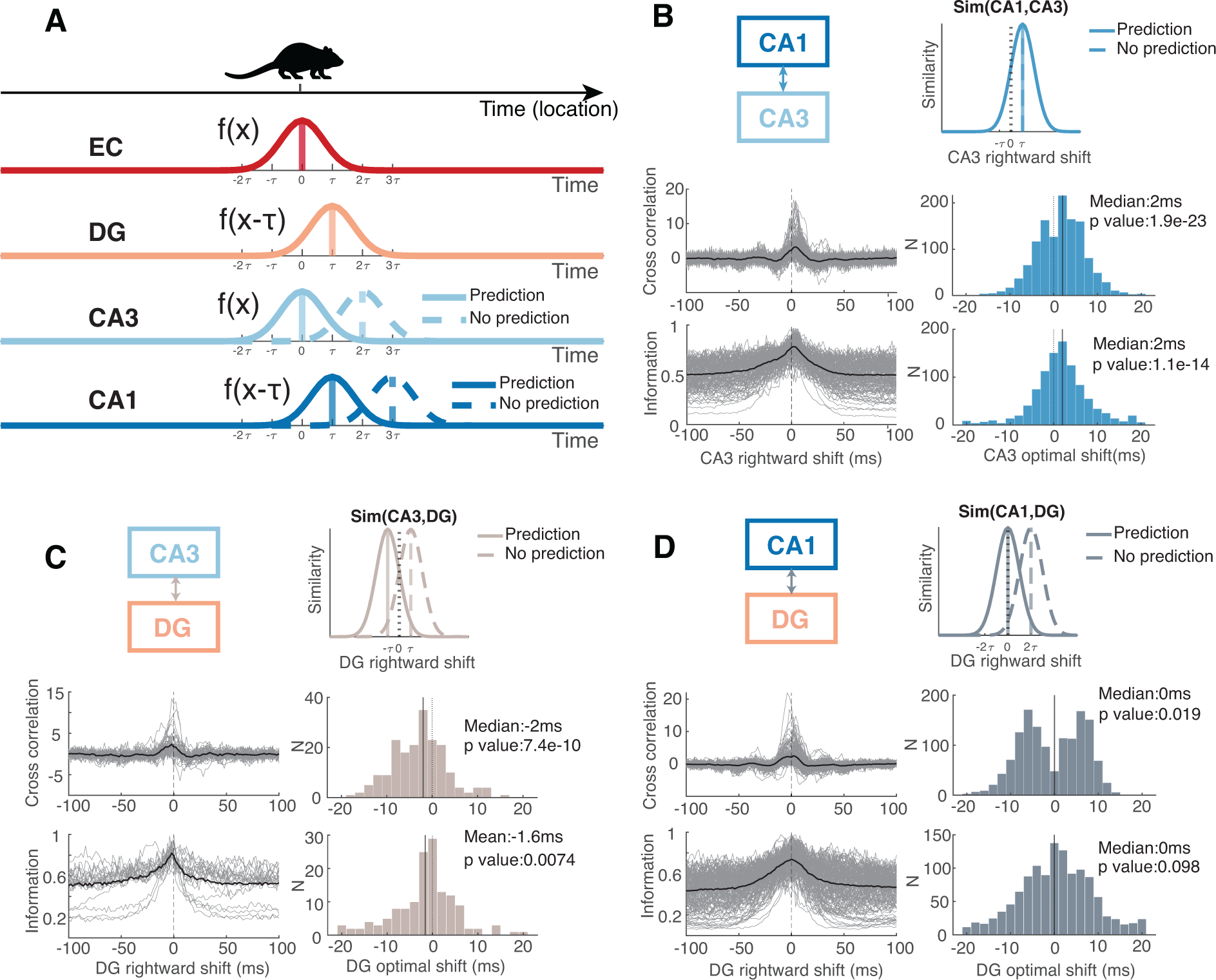
Neural evidence of transmission delay and predicting ahead. (A) Schematics of delayed neural response and hypothesized predicting effect. Assume a rat running with constant velocity (time=location), one representative location sensitive neuron in EC exhibits bell-shaped response curve peaked at *t* = 0. Given there’s no prediction, its direct downstream DG and CA3 neuron will peak at *t* = *τ* and *t* = 2*τ*, respectively. Meanwhile, CA1 would receive mixed signals, delayed by *τ* and 3*τ*, from dual pathways. If there is prediction ahead, CA3 would instead peak at *t* = 0 and CA1 would only respond to signals peaked at *t* = *τ*. (B) Spike coupling from CA3 to CA1. Top: schematics of spike train similarity with respect to CA3 neural activity shifts. Positive shift means shifting CA3 spike train towards the right and then computing its similarity with the unshifted CA1 spike train. Middle (Bottom): Left: Traces of corrected cross correlogram (Mutual information) from an example session. Each gray trace represents the prediction from a population of CA3 neurons to one CA1 neuron. The solid black trace is the average across all CA1 neurons in the session. Right: Histogram of optimal shift, where similarity measure peaks, pooled across 12 recording sessions. (p-value: t-test of population mean equals to zero) (C) Spike coupling from DG to CA3. (D) Spike coupling from DG to CA1. DG is synchronized with CA1 while CA3 leads DG by 2 ms. This confirms the

Alternatively, if, according to our hypothesis, CA3 is predicting future signals to match the signal arrived from the direct pathway, CA3 and CA1 would have response curves of *f* (*x*) and *f* (*x − τ*), respectively (solid lines in Fig. 2A), given similar interregional delays. Although the recordings are unlikely to be from directly connected neurons, evaluating the similarity measures between distributions of temporally shifted neural activities should reveal interregional spike coupling properties (Fig. 2BCD, upper panel).

According to our hypothesis, CA3 spike trains should couple tightly with leftward-shifted DG spike train (Fig. 2C, upper panel), indicating that CA3 firing leads DG. This suggests that CA3 is predicting ahead since it is anatomically downstream from the DG. For both the prediction and non-prediction scenarios, CA1 activity should always follow CA3 by one synaptic delay (Fig. 2B, upper panel). Moreover, despite the challenges associated with measuring CA1-DG coupling due to their lack of direct connection, similarity measures for CA1 and DG spike trains for the prediction scenario are expected to reach a maximum at approximately zero delay since both of these areas are one synapse away from the EC. A peak at zero signifies information synchrony between these two regions and would highlight the predominance of signals delayed by *τ* in CA1 (see Fig. 2D, upper panel).

We used the visual encoding neuropixel dataset from the Allen Brain Observatory (Siegle et al., 2021). This dataset contains simultaneous recordings of neural spikes at a sampled at 30 kHz in DG, CA3 and CA1 from mice performing passive visual perception of natural movies, i.e. sequences of natural images (Methods). The high temporal precision enabled us to investigate spiking timing accuracy on a millisecond time scale.

Following the methods in Siegle et al. (Siegle et al., 2021), we calculated the jitter-corrected cross correlagram (CCG) of spike trains between pairs of subregions over all stimulus conditions and plotted the distribution of optimal shifts where CCG peaks (Fig. 2BCD) (Methods). To access higher-order statistical relationships, we also calculated the mutual information between the shifted spike trains since we are interested in the amount of delay in the information transmitted by the spike trains. When we used shifted CA3 spike trains to predict unshifted CA1 spike train in Fig. 2B, they coupled most strongly when CA1 was shifted to the right. Thus, unsurprisingly, CA3 was ahead of CA1 activity by 2 millisecond, which matched the previously reported synaptic delay (Leung et al., 1995; Sabatini & Regehr, 1999), validating our approaches to calculate synaptic coupling at the precision of milliseconds.

In Fig. 2C, we compared the unshifted CA3 spike train with shifted DG spike trains. We found that both similarity measures peaked when DG shifted significantly towards the left by a median of 2 millisecond. This directly supports the CA3 predictive ahead hypothesis as explained above. In Fig. 2D, we compared the unshifted CA1 spike train with shifted DG spike trains. From the mutual information analysis, the distribution of optimal shifts is approximately a normal distribution with median value of zero. This means that neurons in CA1, despite some randomness, are synchronized with those in DG assessed by mutual information. They were both delayed by one synapse with respect to EC. The identification of a leftward peak (CA1 preceding DG) in the cross-correlation analysis and a synchronized peak in the mutual information analysis strongly supports the predictive-ahead hypothesis.

The distribution of optimal shifts between CA1 and DG from cross-correlogram analysis was bimodal (Fig. 2D). We acknowledge that comparing the correlated activity in CA1 and DG may be problematic, considering their lack of a direct connection and the inherent difficulty in recovering coupling at the millisecond level. However, the leftward peak could only appear with the predictive component in the circuit because otherwise CA1 response from the indirect pathway is always going to lag behind DG response. The presence of a rightward peak in the cross-correlation analysis could be a consequence of fast oscillations in the 100-200 Hz range in LFP recordings in CA1. Fast oscillations were not observed in DG on the same probe (see Fig. S2). An oscillation could induce the bimodal peaks observed in Fig. 2D. The mutual information analysis, which is more robust to firing rate fluctuations, was not bimodal, partially supporting this hypothesis.

### Explaining Observed Spike Coupling with a Predictive Recurrent Neural Network

We developed a predictive recurrent autoencoder model of the hippocampus to compare the time delays in the model to those observed between CA3 and DG. In Fig. 3A, we replicated the temporal relationships identified in the preceding section. Notably, the recurrent units in CA3 encode information at *t* + 2 due to the predict-ahead training. Dashed lines represent delay operations, while solid lines signify network computations governed by the equations in Eq. 1. The input signal *x* originates from the entorhinal cortex (EC) and DG, with DG serving solely as a delay operator in our model. The recurrent signal *h* models CA3 activities, and the CA1 response is computed as a concatenation of prediction errors and predictions (*o*). The dynamics of CA3 and CA1 activities will be explored in subsequent sessions.

**Figure 3:**
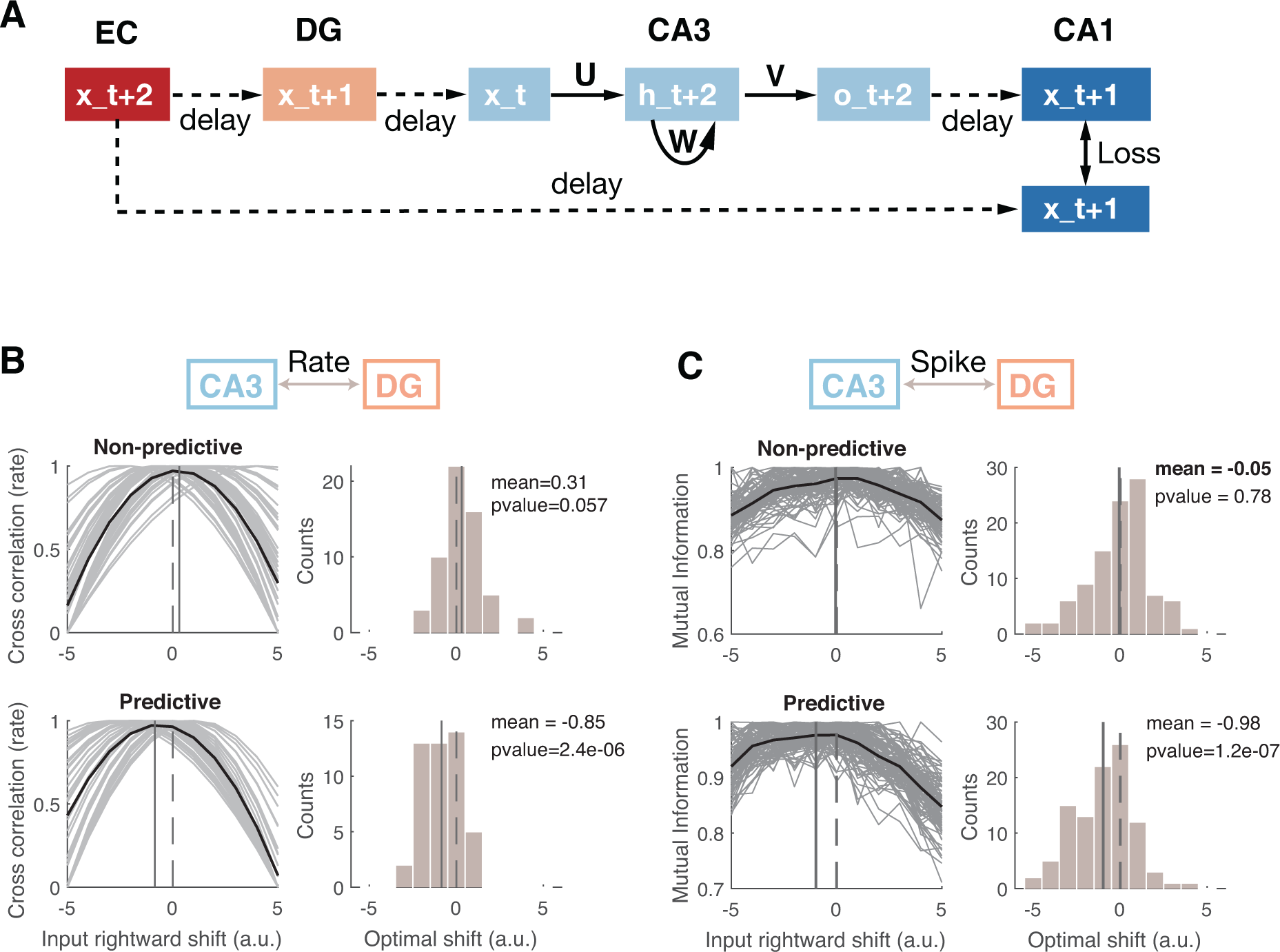
A predictive RNN explains observed statistics. (A) Summary of the temporal relationship observed in Fig.2. The time stamps are labeled from the perspective of CA3 input. Dashed and solid lines indicate the delay operation and weighted computation separately. We believe prediction happened through the recurrent weight *W*, forming a conventional recurrent neural network as described by the equations. EC/DG, CA3 and CA3 function as the input, recurrent and output layer separately. To enforce the recurrent units to predict ahead, we adopted a predictive loss function. (B) Rate coupling from DG to CA3, equivalently, cross correlation analysis between *x_t_* and *h_t_* for non-predictive networks (first row) and predictive networks (second row). In a predictive network, recurrent units correspond better with *x_t_*_+1_. (C) Mutual information analysis of spikes from DG to CA3. Spikes are generated through a Poisson process with the rate given by the trained networks and Δ*t* = 0.02 sec.

To train the model for predicting its next-step input *x*, we adjusted the recurrent weight *W* using backpropagation through time (BPTT) (Bryson, 1961; Werbos, 1990; Rumelhart et al., 1986) over a predictive loss function (Eq. 1). For comparison, we also trained the network using a non-predictive loss function. In the later section, we show that the error signal leads to comparable learning performance with biologically plausible learning rules.

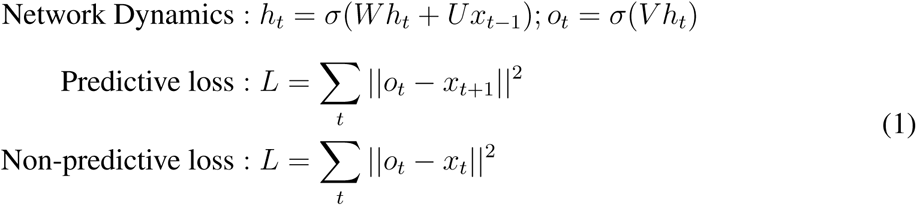

where inputs *x_t_* project to the hidden units *h_t_* with weights *U*. The hidden units are connected with recurrent weights *W*, and project to the outputs *o_t_* through weights *V*. We tasked both networks with learning a Bell-shaped pulse spanning 100 time steps, centered at *t* = 50, and subsequently calculated the cross-correlation of *x* and *h*, as illustrated in Fig. 3B. In the network trained using a predictive loss function, the recurrent units exhibited a notable leftward shift compared to the input (bottom panels). This shift was not observed in the control network trained using the non-predictive loss function. We then used the firing rates derived from the artificial networks to generate Poisson spikes and performed cross-mutual information analysis on the simulated spike trains. The same results were found in the spiking network model as those obtained in the rate-based model (Fig. 3C).

### Facilitating Learning of Internal Models through Sequence Prediction

Our first test of the predictive recurrent network model was to explain place cell dynamics. We first simulated a rat running along a circular track with constant velocity (where time and location are equivalent). All neurons in the recurrent layer (CA3) received location specific bell-shaped input activity (*x_t_*) representing their respective place fields on the track (Fig. 4A, left panel). A new environment was modeled as a random shuffle of place fields (Fig. 4A, right panel). The network was trained on Env1 and Env2 using the predictive loss function.

**Figure 4:**
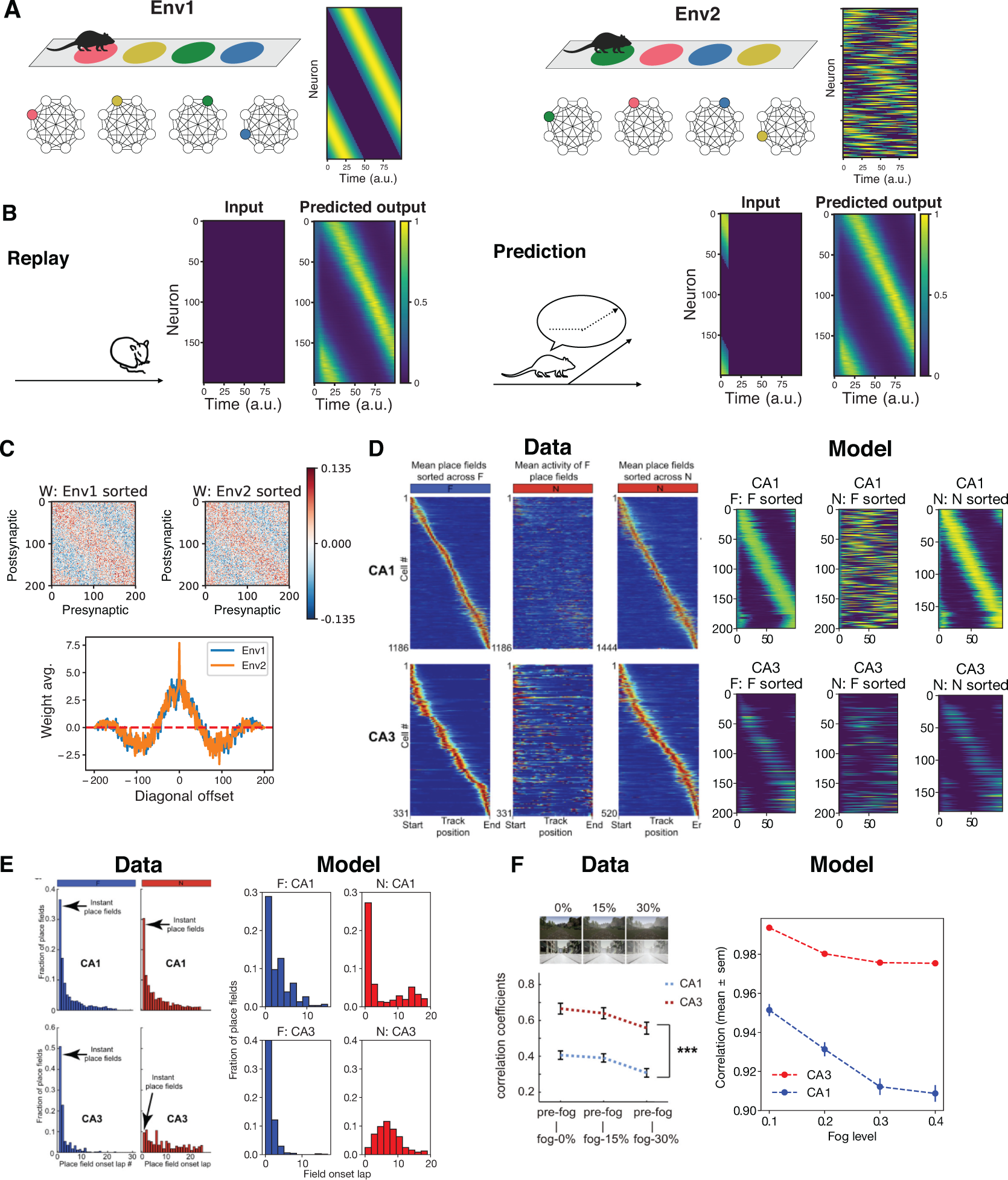
CA1 error encoding neurons facilitate the learning of internal model and explain distinct CA1 and CA3 place field dynamics during remapping. (A) Input matrices (*x_t_*) used during remapping. With the equivalence of time and location, each row represents a bell-shaped location specific input current. Env: environment. A second environment was modeled as the complete shuffling of the first familiar environment (B) Replay: Given low magnitude random input simulating spontaneous activities, a predictive recurrent autoencoder output its previously remembered pattern. Prediction: given input of the first 10 time steps, the network performed pattern completion. (C) Upper: The trained recurrent weight matrix sorted by the activation order of Env1 or Env2. Note similar entries on the diagonals; Lower: Average value of the matrix diagonals offset by the index shown on the horizontal axis. The approximate kernel here looks like a Mexican hat, pointing to the existence of line attractor dynamics. (D) During remapping, different ensembles of neurons are activated in both CA1 and CA3 populations. Place fields of CA1 (first row) and CA3 (second row) neurons in familiar (F) or novel (N) environment sorted by activation order. (Data from Dong et al. (2021) Fig.2). (E) Histogram of CA1 (first row) and CA3 (second row) place field onset laps in F (blue) or N (red) environments. Both regions showed instant onset place fields in F but only CA1 neurons responded instantaneously in N. (Data from Dong et al. (2021) Fig.2). (F) Correlation of CA1 and CA3 place fields between F environment and its noisy/foggy version. See Table 2 For details about the network and training. Data from Shin et al. (2022) Fig. 5.

The trained models successfully reproduced the input sequences and exhibited replay and prediction (Fig. 4B). Replay refers to the re-activation of place cells in the same order as they would during active exploring. Typically, this occurs when the animal is in a state of sleep or immobility, meaning that the simulated agent is not receiving any external sensory inputs. When low magnitude random noise was used to drive the trained network, it randomly reproduced one of the learned sequences (Fig. 4B, left and Fig.S3).

Place cells in the model also showed predictive activities that have been reported for cells that are activated before making turns and code for possible future locations (Ólafsdóttir et al., 2018). This could be a consequence of pattern completion by the recurrent network model. To demonstrate this, following a partial input sequence to the network, it completed the remaining sequences (4B, right).

This is strong evidence for line attractor dynamics (Eliasmith, 2007) in the network. We reordered the learned recurrent weight matrix based on the activation order of the hidden units in either Env1 or Env2 (Fig. 4C, upper). The reordered matrix has an approximately symmetric Toeplitz form resembling a one-dimensional chain of neurons, each connected to its nearest neighbors with positive values and more distant neurons with negative values (Fig. 4C, lower). The mathematical significance of Toeplitz connectivity is elaborated in the Discussion.

The trained network also exhibited phase precession, which occurs in recordings where the timing of place cell firing with respect to the phase of the oscillatory population activity becomes progressively earlier when traversing a place field (O’Keefe & Recce, 1993). To generate biologically relevant action potentials, we transferred the weights from a trained recurrent weight to a network of leaky-integrate-fire (LIF) neurons following the procedure described in (Kim et al., 2019) and recorded the emitted spikes (Fig. S4A). Oscillatory activity was artificially enforced by injecting 8 Hz inhibitory currents, mimicking oscillatory inputs onto inhibitory neurons originating in the septal nucleus. Spike phases were calculated and plotted against their relative location to place field centers. Analysis of the LIF neurons during simulated running on the track exhibited precession of the spike timing (Fig. S4B) similar to phase precession recorded from neurons *in vivo* (See Fig. 1 in (Tsodyks et al., 1996)).

### CA1 error encoding neurons explain distinct CA1 and CA3 place field dynamics

Although differences in the encoding properties of place cells in CA3 and CA1 are well known (Leutgeb et al., 2004), they have been overlooked in most hippocampal models. In our model, CA3 stores an internal model of the world, while CA1 not only inherits CA3 output but also simultaneously encodes prediction error. Supporting evidence for error-encoding neurons involves earlier experimental observations that CA1 neurons respond more than neurons in other regions to unexpected signals (Kumaran & Maguire, 2006; Knight, 1996; Duncan et al., 2012) and that CA1 place fields decay slowly in familiar environments (Fig.S1). At the same time, acute silencing of CA3 drastically affects CA1 response (Davoudi & Foster, 2019), suggesting CA1 also largely inherits CA3 output. In a later section we present a biologically plausible predictive learning algorithm for CA3 based on local prediction error.

To model remapping from a familiar environment (F) to a novel environment (N), we instructed a network that had already memorized Env1 to optimize towards Env2. Neural responses in CA3 and CA1 were recorded as described above and sorted based on experimental observations (Fig. 4D). For both CA1 and CA3 neurons, distinct ensembles of place cells were activated in the two environments, consistent with previous experimental data (Dong et al., 2021). Importantly, in Fig. 4E (right panels), upon the switch to the novel environment, CA1 place cells emerged more rapidly, as error information initially reflected the structure of the novel environment. Over time, CA1 place cells transitioned to representing CA3 output. In contrast, CA3 place cells emerged more slowly, suggesting that the internal representation might require multiple traversals to learn. The rapid neural response upon exposure to a novel environment aligns with the well-established place cell property called one-shot learning (Priestley et al., 2022), the mechanism of which is still debated. We propose here that this may be attributed to the coding of error signals rather than an abrupt increase in weights. Abruptly modifying synaptic weights to a large population of neurons (Bittner et al., 2017) could pose risks to system stability if not tightly regulated. In this study, we present an alternative explanation for apparent one-shot learning: neural activity immediately increases in a new environment because CA1 reflects one-shot error, not one-shot learning.

Next, we modeled place cell remapping back to a familiar environment by optimizing towards a noisy version of Env1 (Fig. 4E, left panels). Consistent with experimental data (Dong et al., 2021), both regions exhibited instant place fields, as the internal representation stored in the network matched the familiar environment. We also examined the relationship between the correlation of neural activities and the noise level in the Env1 environment (i.e., fog percentage in the data panel); noise decorrelated the representation in both CA3 and CA1, but CA3 was more robust to the presence of noise (Fig. 4F).

### Learning future sequences promotes interpretable latent representation

We next simulated a rat in a random foraging task consisting of straight trajectories and random turns in a square arena (Fig. 5A). Input was handcrafted as a nonlinear mixture of body direction, world direction, path integrated distance and distance to the closest wall (Methods) following the approach described in (Benna & Fusi, 2021). The selection of these input dimensions was based on the existence of head direction cells, path-integration signals and border cells in EC. Any one of these inputs alone is not able to determine the agent’s current location, as evidenced by the low mutual information per second (MI) between input unit activity and location (Fig. 5 C). To enforce a sparse representation in the hidden layer, we added a regularization penalty of unit activity to the MSE loss function (Methods).

**Figure 5:**
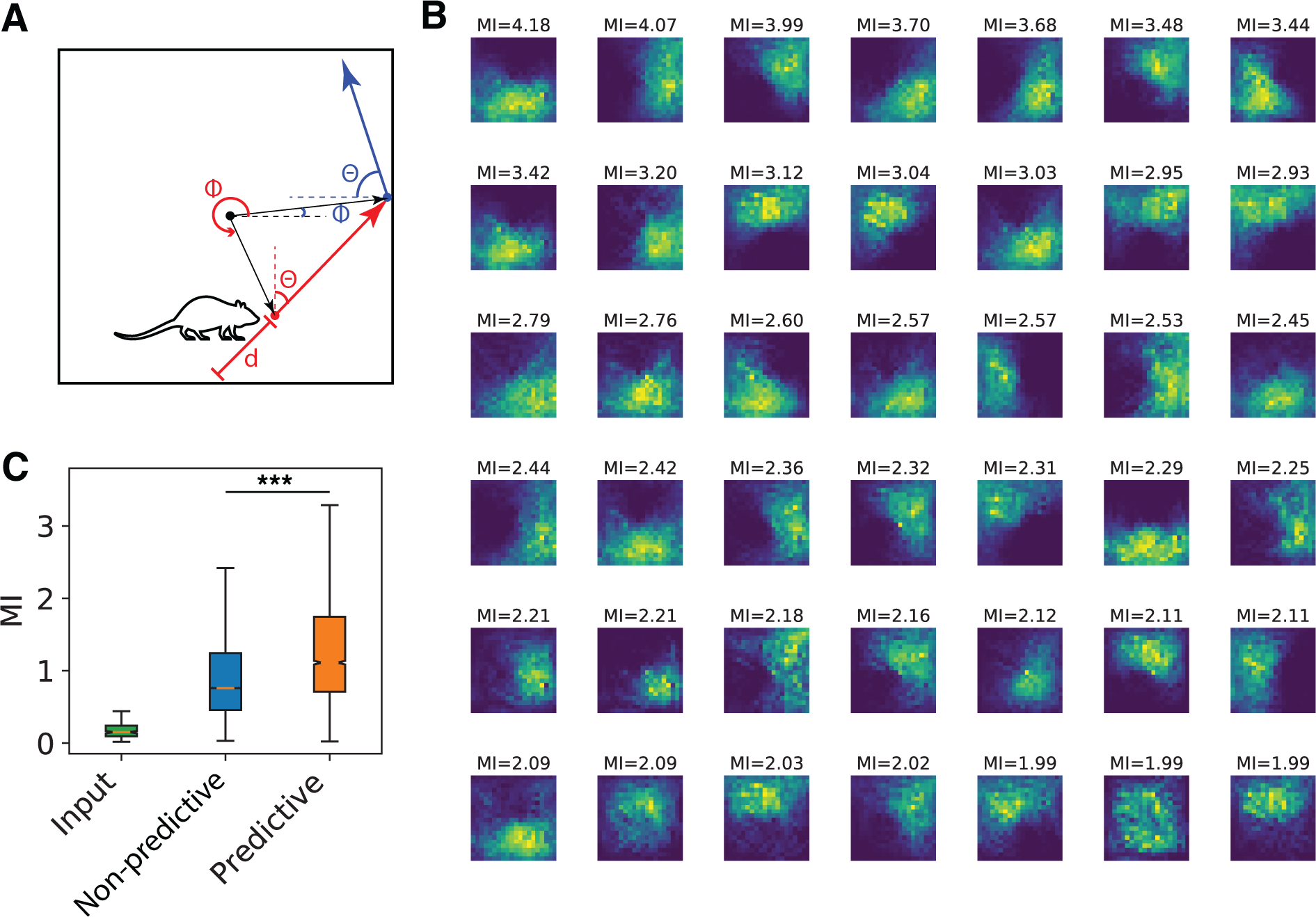
Predictive networks led to more localized representation in a random foraging task. (A) Schematic of foraging task simulation. The agent was running straight in an open arena until it hit a wall and make a random turn. The red and blue straight arrows indicate two straight trajectories. Network input defined as a random nonlinear mixture of body direction with respect to the norm of the last hit wall (*θ*), world direction with respect to east (*ϕ*), path integrated distance (*d*). (B) Place fields of the recurrent units (CA3 neurons) after training. Extent of localization was quantified by mutual information (MI) rate per second between firing rate and location. (C) MI of the input units and recurrent units in 10 networks trained by either the current loss function as a control or the predictive loss function. Recurrent units trained by the predictive loss function showed significantly higher MI (t-test, p*<* 0.001).

After training using the predictive loss function, hidden units showed spatially localized representations similar to place fields. (Fig. 5B). In comparison to networks trained with the non-predictive loss function control, hidden units in networks trained with the predictive loss function displayed significantly higher MI (Fig. 5C). Both networks contributed to the extraction of locations as hidden units have much higher MI compared to the original input signal. This indicates predicting ahead is beneficial for extracting location information from upstream inputs. The same results were found when the regularization strength was varied (Fig. S6), suggesting that a temporal predictive loss function consistently aids in forming a localized representation.

To investigate the potential of the predictive loss function for achieving representational learning—specifically, forming sequential representations from high-dimensional sensory inputs—we organized image sequences using MNIST handwritten digits. We first trained the network to predict sequences of increasing MNIST digits (Fig. 6). Multiple batches of randomly sampled images were temporally ordered based on their labels. We extracted the first 68 principal components (PCA) of the images, which account for 86% variance, as the input to keep the minimal network structure. Image reconstruction was based on the network output and inverse transformation of PCA. After training, the network continued to complete the sequences when the input was stopped after digit 3 (Fig. 6C). Predictive completion was not observed in control models trained with the non-predictive loss function (Fig. S5).

**Figure 6:**
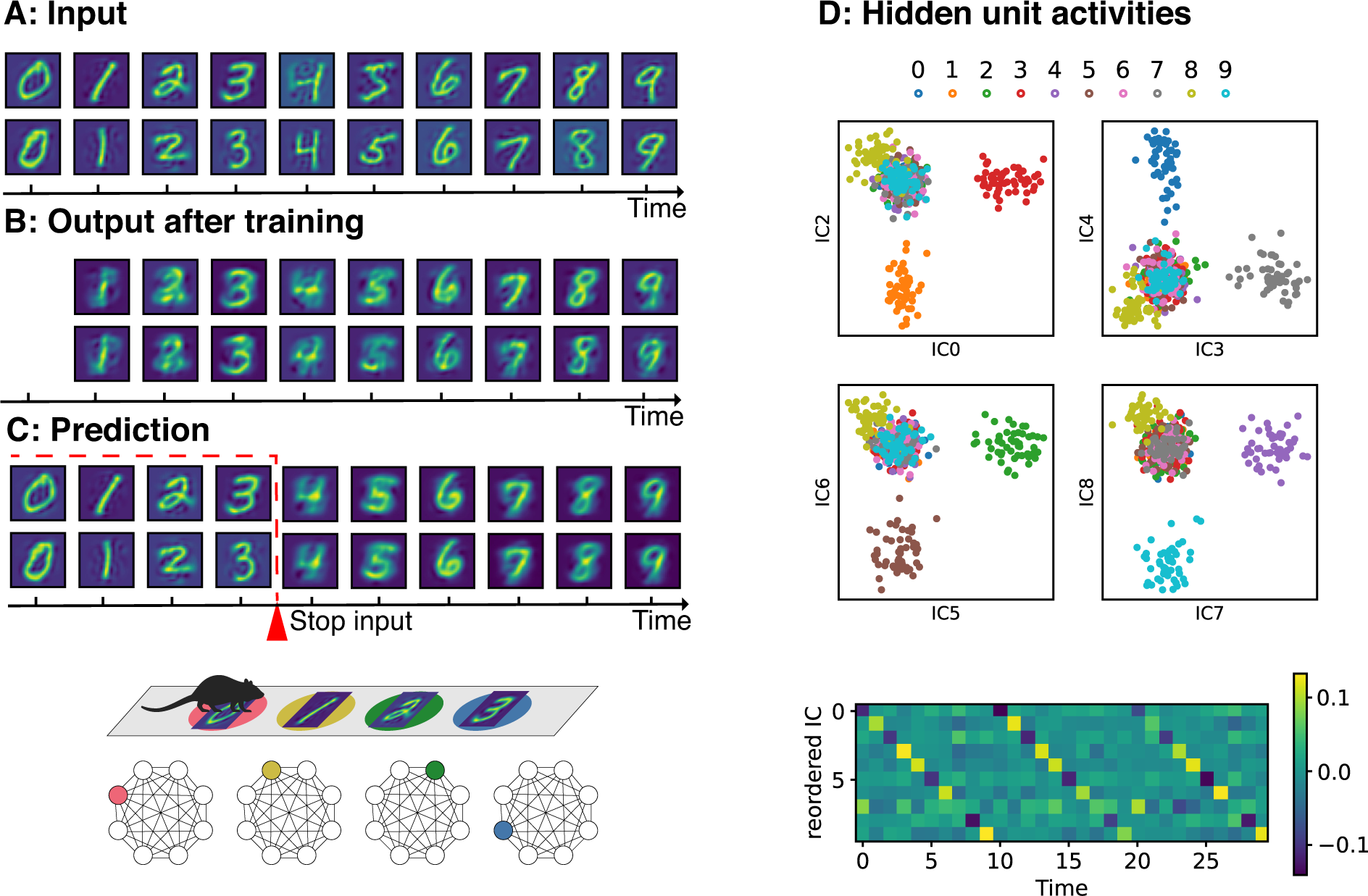
A predictive networks compress visual cues into sequential activation of hippocampal neurons. (A) Top: Input organized as repetitive sequences of MNIST images from 0 to 9. Two sequences show individual differences. (B) Trained network initiated with the first input could reconstruct the subsequent generic digits. (C) When the input was stopped after digit 3, the network continued to predict the rest of the digits. Input digits were plotted before the red dashed line while predictions were plotted afterwards. (D) Top: Independent components (IC) of hidden unit activity. Most ICs represent one class (color-coded); Bottom: Reordered IC activities over time. Note the sequential activation in 10 time steps (one cycle). ICA does not extract a unique sign, so diagonal entries can have both large positive and negative values.

Interestingly, the trained network not only constructed a generative model to predict sequences but also automatically clustered the digits according to their labels. In Fig. 6D, we plotted the independent components (IC) of the hidden unit activity for each digit and colored the digits according to their labels. For the top 10 ICs (sorted by the contribution of demixing matrix), approximately each component represents one group of digits.

This interpretable representation was achieved without explicitly defined labels. The sequential activation of ICs along the time axis (Fig. 6D, bottom panel) mimics the sequential activation of place cells in the linear transformed space. Imagine a rat running on tiles of MNIST patterns: Fig. 6 shows how the interaction between cortex and hippocampus transforms the complicated sensory input into sequential activation of hippocampus neurons as reported in numerous experimental studies.

### Error Neurons Facilitate a Biologically Plausible Learning Algorithm

We devised a predictive recirculation learning algorithm for a predictive autoencoder, consisting of a set of three local learning rules for the input weights (*U*), recurrent weights (*W*), and output weights (*V*), (Eq. 2). These learning rules approximate the gradient of a predictive mean square error loss under certain assumptions (see Methods for derivations).

The precise gradient of the output weight (Δ*V* in Eq. 2) can be directly assessed as Hebbian learning between error encoding neurons in CA1 (*δx*) and the recurrent neurons in CA3 (*h*).

The exact gradients of the input and recurrent weights pose challenges to achieving locality in both time and space. This temporal dependence was mitigated by truncating the temporal gradient beyond the current time step. Additionally, to preserve spatial locality, we avoided backpropagating errors by using recirculation. This strategy, inspired by the original recirculation algorithm proposed by Hinton & McClelland (1987) for a three-layer feedforward autoencoder, facilitates local learning for both input and output weights by feeding back reconstructed inputs to the encoder (Eq. 4). As the authors of the recirculation algorithm noted, the input weights converge approximately to the transpose of the output weight (*U* = *V ^T^*). We confirmed that during learning (Eq. 2), the matrix entries in *U* and *V ^T^* in our predictive autoencoder also converged: Predictive recirculation learning effectively drove the weights from random initialization to approximate transposition (Fig. S7). In the hippocampus, we propose that this recirculation process could be implemented through the feedback projections from CA1 to EC (Fig. 7A).

**Figure 7:**
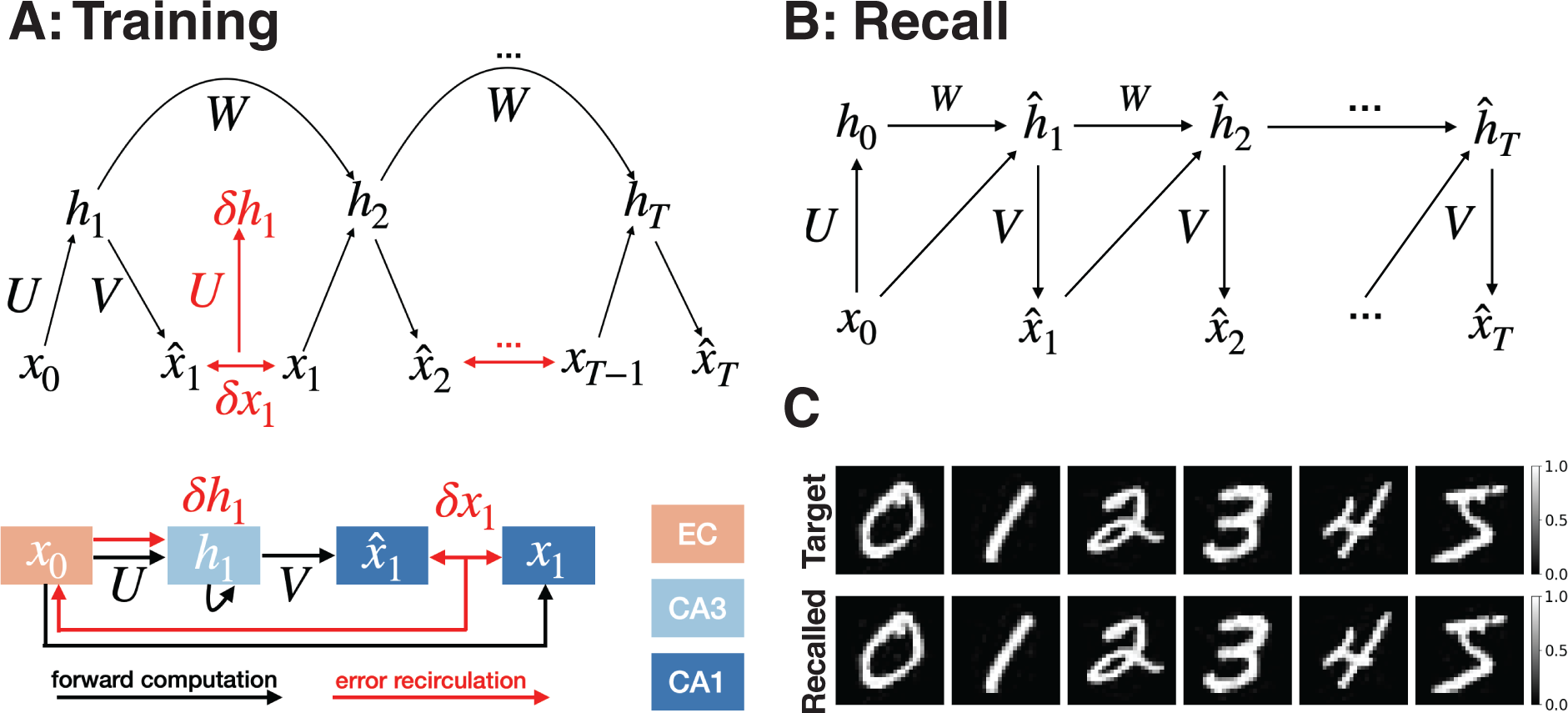
Predictive recurrent network that can learn and playback sequences of images using biologically plausible local learning rules. (A) During training, inputs *x_t_* project to the hidden units *h_t_* in the recurrent network. Three local learning algorithms update the weights *U* from the inputs to the hidden units, the weights *V* that feedback to the inputs, and the weights *W* between the hidden units in the recurrent network (Methods). All three weight updates can be computed by prediction error *δx* and its recirculated error *δh* = *Uδx* through the feedback pathway from CA1 to EC. (B) During recall, an initial input *x*_0_ generates a sequence of outputs from the hidden layer. (C) Example of a network trained by local learning algorithms on an MNIST sequence of handwritten digits (Target). The output following the 0 input without further input replays the sequence in the trained order (Recalled). See Table 2 For details about the network and training hyperparameters.

In Fig. 7, we trained a network using predictive recirculation algorithm to reproduce and recall a sequence of MNIST handwritten digits.

#### Predictive recirculation learning algorithm

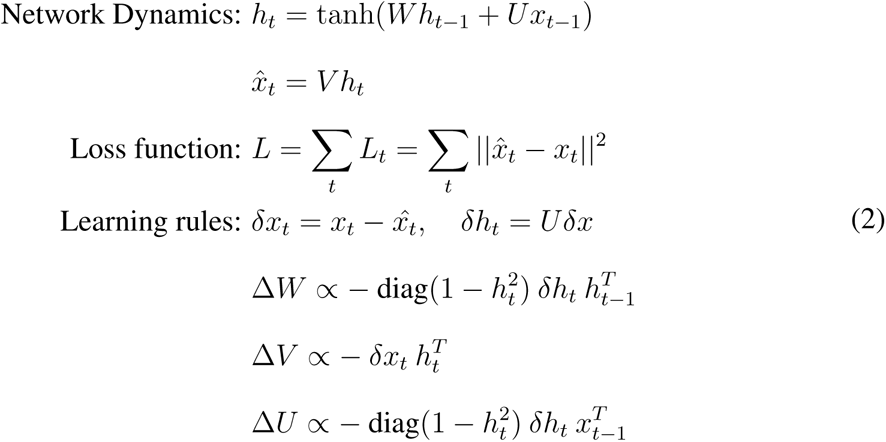

where diag(*y*) is a diagonal matrix with its diagonal equal to the vector argument *y*. A derivation of these learning algorithms is given in the Methods.

## Discussion

The temporal predictive coding framework proposed here to account for sequence memory and representation learning was inspired by the anatomy of the hippocampus, validated by neural recordings, and successfully replicated a variety of experimental observations.

This theoretical framework makes several experimental predictions: First, the ablation of the direct pathway is expected to suppress the formation of new place fields upon remapping. This is partially supported by findings from Grienberger et al (Grienberger & Magee, 2022), where the optogenetic inhibition of EC3 input activity led to a significant reduction in experience-dependent shaping of CA1 representations. Second, our analysis suggests that the significance of CA3 predictions may grow slowly during the early stages of sequential learning, requiring multiple epochs of training to achieve accurate prediction. Third, our model makes detailed predictions that could be tested with simultaneous long-term recordings from CA1, CA3, and EC recordings before and after learning a sequential task. Analysis of time course of coupling between different regions would reveal the amount of prediction and how it changes over time.

We found that training on sequences yielded a recurrent network with a sparse Toeplitz form that can store multiple sequences. Toeplitz matricies have a diagonal structure that performs a matrix temporal convolution. Toeplitz matricies also support traveling waves, which speed up learning of sequential tasks by two orders of magnitude and over much longer time scales (Keller et al., 2023). The Toeplitz convolutional kernel underlies moving-bump line attractor dynamics in recurrent neural networks, including the dynamics of network models for compass cells in rodents (Zhang, 1996), neural integrators, and other neural systems (Gardner et al., 2022; Khona & Fiete, 2022). A connectomic analysis of the rodent area CA3 could potentially confirm the predicted Toeplitz connectivity, providing further validation for our proposed model.

Predictive loss functions are routinely used in state-space models, such as model-based control and Bayesian filtering. By predicting the future observations, generative models are continually updated to make more accurate predictions (Isomura & Toyoizumi, 2021). This approach has recently been incorporated in several model-based artificial intelligence systems (Recanatesi et al., 2021; Lotter et al., 2016). The performance of these systems is more robust and has superior generalizability when there is an internal model of the system. Our predictive model-based approach to the hippocampus has similar advantages and is supported by evidence from neural recordings.

Recanatesi et al. (Recanatesi et al., 2021) also explored predictive network models with both state and action as inputs to predict the state on the next time step. They demonstrated that representation compression, including localized representation from high-dimensional input, could be achieved through the addition of action signals. However, the evidence for action signal encoding in the entorhinal cortex, the main input to the hippocampus, is minimal. Without the action input, we also observed compression of redundant representation in Fig. 5 and Fig. 6 as long as the input signal is redundant, indicating that this is a robust computational advantage inherent in having a predictive loss function. Our predictive model without action aligns closely to a one-step successor representation (Fang et al., 2023).

Previous studies have also explored the anatomy and firing properties of hippocampal neurons (Basu & Siegelbaum, 2015; Klausberger, 2009), based on a wide range of theoretical and computation principles (Stachenfeld et al., 2017; Whittington et al., 2020; McNamee et al., 2021; Benna & Fusi, 2021). Stachenfeld et al. (Stachenfeld et al., 2017) modeled the activities of place cells as the successor representation in reinforcement learning; Whittington et al. (Whittington et al., 2020) proposed that the hippocampus is a statistical machine for inferring structural properties from observations. Our model is more mechanistic and based on the analysis of neural recordings from hippocampal subregions. The simplicity and minimal realization of our model could also serve as a critical building block for statistical inference.

Representing prediction errors in some CA1 neurons could not only facilitate the learning of the internal model stored in the recurrent CA3 network, but could also regulate the release of dopamine in novelty-dependent firing of cells in ventral tegmental area (VTA) through subiculum, accumbens, and ventral pallidum (Lisman & Grace, 2005). This could explain why novelty detection is an essential function of the hippocampus. Temporal prediction error has already been established for learning sequences of actions in the basal ganglia to obtain future rewards (Watabe-Uchida et al., 2017). If temporal predictive coding principles for learning sequences are also found in the cortex, as suggested in Fig. 8, then predicting the next input in every cortical area may be an important design principle for human cortical function.

**Figure 8:**
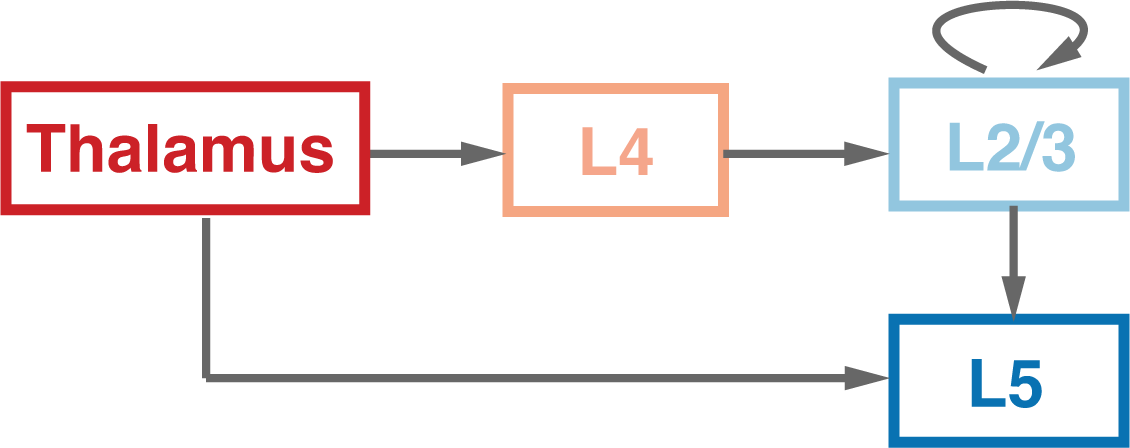
Universality of the circuit for computing temporal prediction error in the cortex. Signals from thalamus reach cortex Layer 5 through two different pathways: the direct pathway (Gökçe et al., 2016) and the indirect pathway via Layer 4, recurrent Layers 2/3 (Markov et al., 2013). This cortical circuit resembles the pathways in the hippocampus (Fig. 1): The small stellate cells in Layer 4 may have the same preprocessing function as DG granule cells; recurrent Layers 2/3 learns sequences by making temporal predictions; Pyramidal neurons in Layer 5 compute the temporal prediction error between these direct and indirect pathways, and propagated globally to subcortical structures and through Layer 6 to the reticular nucleus of the thalamus. A hierarchy of sequences learned by temporal prediction may be computed in the neocortex (Haeusler & Wolfgang, 2007) since the canonical circuit is similar throughout.

Self-supervised models, such as variational autoencoders (Kingma & Welling, 2019), unsupervised Boltzmann Machines (Ackley et al., 1985; Hinton et al., 1986) and their many variants, have avoided labor-intensive supervised input labeling. The recent success of self-supervised transformers like GPT were trained by predicting the next word in a sentence (Vaswani et al., 2017; Sejnowski, 2023). These sophisticated models implicitly learn semantic representations. In the same way, our predictive network achieves representation learning of sequences without needing sophisticated statistical priors or explicitly defined representation modules. Its minimal model structure facilitates detailed investigation and interpretation. Future work could focus on scaling up the network or adding preprocessing modules to handle more realistic problems such as the semantic segmentation of video clips.

There is an analogy between the superficial and deep layers of the six-layered neocortex with areas CA3 and CA1 of the hippocampus, respectively. This is illustrated in Fig. 8. Upon comparison with Fig. 1, parallels emerge between the indirect pathway through the DG in the hippocampus, projecting to CA3, and the thalamic inputs to layer 4 of the neocortex, projecting to layers 2/3. Similarly, the direct pathway from the EC to area CA1 in the hippocampus corresponds in the cortex to direct inputs from the thalamus to layer 5.

As in the dentate gyrus, layer 4 neurons are small and numerous, creating input representation that separate similar patterns; neurons in layers 2/3 form a highly recurrent network, similar to that in CA3; neurons in layer 5 are output neurons, like CA1 neurons. This similarity has been noted by others.^1^ We go further and suggest that all cortical areas, as well as the hippocampus, may be predictive autoencoders.

In this cortical model, the recurrent network in layers 2/3 is trained as a predictive autoencoder to remember sequences of inputs arising from the thalamus, which, like the EC in the hippocampal model, serves as the input and output of the autoencoder. The latent sequences are further compressed in the downstream areas, becoming more abstract while ascending the hierarchy. The EC in Fig. 8 is at the top of converging hierarchies of cortical areas, each recapitulating the same architecture, forming stacks of predictive autoencoders.

This generalization of our hippocampal model could be tested by recording simultaneously from the thalamus, layer 4, layers 2/3 and layer 5 and analyzing the spikes the same way we analyzed recordings from the corresponding areas of the hippocampus, DG, CA3 and CA1.

If temporal predictive coding principles for learning sequences are found in the hippocampus and the cortex, as suggested in Fig. 8, then predicting the next input in every cortical area may be an important design principle for human cortical function. Predictive learning using conserved circuits could underlie the robustness and flexibility of human intelligence. Transformers in large language models achieve remarkable performance with predictive self-supervised learning. Inspired by brains, PredRAE has potential for further improvements in robustly disentangling representations in artificial intelligence and approaching human levels of performance.

Predicting the next input may be an even more general design principle for brains. Reward prediction error has already been established for learning sequences of actions in the basal ganglia to obtain future rewards (Watabe-Uchida et al., 2017). The temporal difference learning algorithm compares the expected reward with received reward on each time step. The computational strength of this principle is illustrated by Generative Pretrained Transformers (GPT) in Large Language Models, which are trained to predict the next word in text (Sejnowski, 2023).

Although we used backpropagation through time as a way to construct networks that learn sequences, we showed that it could potentially be replaced by local learning rules by combining multiple biologically plausible learning algorithms that we call Predictive Recirculation. Our local learning rule computes a temporally truncated version of the gradient computed by back-propagation through time. It might nonetheless be difficult to accumulate gradients for long sequences. It is possible that the pathway from the entorhinal cortex to CA3 might facilitate the learning of longer sequences. The addition of grid cells from EC might also make it easier to learn long sequences (Chandra et al., 2023). CA3 may also behave like a reservoir (Jaeger & Haas, 2004), generating a wide range of time varying signals and the prediction error signal could be used to select inputs and weight them to reduce the prediction error. We will pursue these possibilities in a subsequent study.

## Funding

DARPA W911NF1820, ONR N00014-23-1-2069, Swartz Foundation.

## Authors contributions

Y.C., H.Z., T.S. conceptualized the study, wrote and revised the manuscript; Y.C, simulated the computational models, and Y.C and H.Z analyzed the experimental data. We thank Homero Esmeraldo for early exploration of biologically plausible learning algorithms.

## Competing interests

No competing interests to be declared.

## Data and materials availability

The Allen NeuroPixel and CRCNS (hc-6) datasets used in the analysis is publicly available through their website. All neural data analysis scripts and BPTT trained models used in this study are available in https://github.com/yschen13/HCPrediction. The local learning rules are available in https://github.com/miacameron/Local_RNN.

## STAR Methods

### Neural evidence of transmission delay and predicting ahead

We used the publicly accessible visual encoding NeuroPixel dataset (Siegle et al., 2021) from the Allen Brain Observatory. Neuropixels were used to simultaneously record the spiking activity of thousands of neurons in mice passively perceiving standard visual stimuli such as drifting gratings, natural scenes, natural movies and et al. We pre-selected recording sessions that involves recordings from DG, CA3 and CA1 in “Functional Connectivity” in WT mice. Very few neurons were recorded from EC. Number of of units being used was summarized in Table 1. For mutual information calculation, only recordings from the natural movie viewing sessions (30 seconds *×* 80 repeats) were used while for cross correlation, recordings from all stimuli sessions were concatenated to increase signal-to-noise ratio.

**Table 1:** Number of simultaneously recorded neurons from DG, CA1 and CA3 in each session.

| Session | CA1 | CA3 | DG |
| --- | --- | --- | --- |
| 766640955 | 170 | 17 | 62 |
| 771160300 | 282 | 48 | 32 |
| 767871931 | 104 | 20 | 32 |
| 768515987 | 102 | 27 | 25 |
| 771990200 | 70 | 6 | 31 |
| 778240327 | 251 | 24 | 42 |
| 778998620 | 133 | 45 | 16 |
| 779839471 | 142 | 25 | 49 |
| 781842082 | 114 | 19 | 28 |
| 793224716 | 159 | 31 | 29 |
| 821695405 | 56 | 17 | 40 |
| 847657808 | 189 | 6 | 70 |

The processed spike train was binned at 2 millisecond (ms). To compute cross-correlogram (CCG) between *N* neurons in region A and *M* neurons in region B, we first calculated jitter-corrected correlograms (Siegle et al., 2021) between *N × M* neuron pairs using a jittering window of 20 millisecond. Jitter-correction was performed by randomly shuffling the spike train within the chosen time window, calculating the jitter-CCG repeatedly for 100 times, and then subtracting its average from the original CCG. Corrected-CCG was further normalized by the geometic firing rate of the neuron pair. In this way, slower time scale correlations, such as the strong theta oscillation in hippocampus or nonstationary trend could be removed and then we could focus on fast time scale neural coupling. To increase signal-to-noise ratio for prediction of one neuron in region A, we used the CCG average of all *M* neurons in region B. Time shifting was performed in region B neurons. For a spike train denoted by *f* (*t*), a positive *τ* shift would lead to a rightward shifted spike train of *f* (*t − τ*). The optimal time shift is defined as the time shift that maximizes the *M* -to-1 averaged cross correlation. We focused on time shifts from −20ms to 20ms as any shifted coupling above this range would be scrambled by jittering.

We compute the mutual information between the spike train of one neuron from region A and the shifted spike trains of 10 neurons from region B. The one-dimensional spike train from region A was treated as a random variable *A*. The latter high-dimensional spike train is treated as a random vector *B* which has 2^10^ states being sampled at different time steps. The mutual information is then calculated as *I*(*A*; *B*) = *H*(*B*) *− H*(*B|A*). For each neuron in A, we compute the information between that neuron and 10 neurons in B and repeat 100 times for different randomly sampled subsets of 10 recorded neurons from B. To show the results for one neuron in A, for each subset of 10 neurons from B, information over time shift is normalized by its maximal value. Then the average of the 100 normalized information curves is taken to reveal the effect of time shift on the mutual information. The optimal time shift is defined as the time shift that maximizes mutual information.

### Analysis of CA1 activity with respect to environment familiarity

Datasets were obtained from http://crcns.org/data-sets/hc/hc-3, contributed by the Buzsáki laboratory at New York University (Mizuseki et al., 2013)(Mizuseki et al., 2014). See http://crcns.org/files/data/hc3/crcns-hc3-processing-flowchart.pdf for more details about experiments, recording and data pre-processing. For rats exploring a 180 × 180-cm box, all sessions that have more than 50 simultaneously recorded CA1 neurons were included for analysis. We excluded neurons that are marked as inhibitory or not identified. For each session, we compute spike rate of the neurons during the last two-thirds of the sessions for stability of the responses.

### Network training

The network was implemented in PyTorch (v1.11.0) and training was performed through stochastic gradient descent of samples split into mini batches with a fixed learning rate, as shown in Table 2. Gradients were calculated with backpropagation through time (BPTT). We used ‘sigmoid’ and ‘tanh’ nonlinearities for the activation of output and recurrent units, unless otherwise mentioned. We stopped training when the process reached the maximum number of epochs or the loss function reached less than 1% of it initial value and did not change more than 0.001% in 10 consecutive iterations. Network structure and related hyperparameters are summarized in Table 2.

**Table 2:** Details of network input and hyperparameters used for simulation. *S*: sample/batch size; *N* : number of input units; *T* : sequence length; *H*: number of hidden units; *L*: loss function; *η*: learning rate.

| Task | $S$ | $N$ | $T$ | $H$ | $L$ | $\eta$ | Total epochs |
| --- | --- | --- | --- | --- | --- | --- | --- |
| Fig. 4 | 1 or 2 | 200 | 100 | 200 | MSE | 0.01 | 50,000 |
| Fig. 5 | 50 | 200 | 100 | 500 | MSE + $\text{Reg}(h_t)$ | 0.01 | 50,000 |
| Fig. 6 | 5 | 68 | 100 | 200 | MSE | 0.01 | 50,000 |
| Fig. 7 | 1 | 68 | 6 | 100 | MSE | 0.0001 | 100,000 |

### Localization

The open arena was simulated as a 2 m *×* 2 m environment. Exploratory trajectory was generated as straight lines of 0.1 m step size until hitting a border. Then a random turnaround angle will be generated to continue exploration. Altogether 5,000 time steps split into 50 samples were used to train the network. Following (Benna & Fusi, 2021) (Supplementary Eq. 1), the location information (path integrated distance, distance to the closest border, world direction and head direction) was randomly and nonlinearly expanded into higher dimensions (*N* = 200) as input and target signal. We switched to ‘ReLU’ nonlinearity for hidden unit activation as we would like to avoid negative responses in terms of place field calculation. To enable a sparse representation, a penalty of hidden unit firing was added to the loss function (Eq.3)

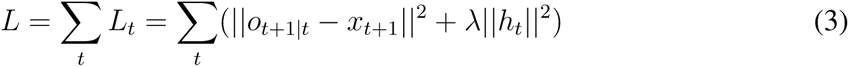

Mutual information of a hidden unit place field was calculated following (Skaggs et al., 1992) as the mutual information between firing rate and the arena location discretized into 25 *×* 25 grids. Specifically, It was calculated as 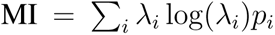 in bits/second where *i* represents location grid, *λ_i_* is the neuron’s firing rate at location grid *i* and *p_i_* is the occupancy probability in grid *i*.

### Learning MNIST sequences

Input was constructed as the top 68 principle components (PC) of the entire MNIST dataset, which explain 87% variance. Input was organized as sequences consisting of 100 time steps, which repeats from digit 0 to digit 9 for 10 times. Five randomly sampled batches of digit images were used for training to predict the next time step PC vector. Independent component analysis (ICA) was performed to reduce the dimension of hidden unit activation from the number of hidden units to the number of chosen ICs (i.e. 10). We manually ordered the ICs by the contribution (column L2 norm) of the converged demixing matrix. For the local learning rule, the input was a single sequence consisting of 6 time steps, where the PC’s were normalized to be between 0 and 1.

### Predictive Recirculation: A biologically plausible learning algorithm

#### Recirculation

The recirculation learning algorithm (Hinton et al., 1986) for a three-layer feedforward autoencoder approximates gradient descent without the need to backpropagate (BP) errors under certain conditions:

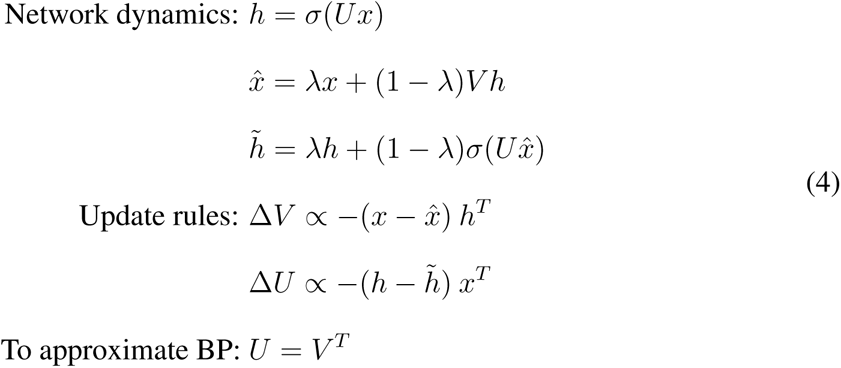

Using this set of learning rules, the symmetry between the input and output weights (up to scaling) is almost guaranteed. A new predictive recirculation learning is derived here based on Eq. 4, and assuming that *U* = *V ^T^*.

#### Output weights

With the dynamics defined in Eq. 2, the exact gradient of the output weight *V* can be obtained using only local information, assisted by error-encoding neurons:

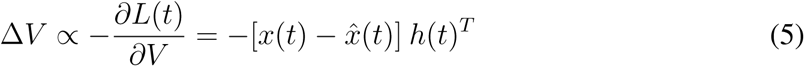

#### Input weights

The exact gradient of input weight (*U*) is given by:

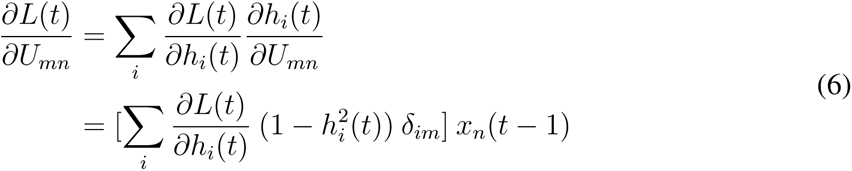

where in our simulations we chose *σ* = tanh, and *σ^′^* = 1 *−* tanh^2^.

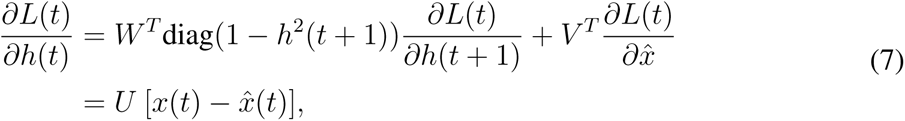

where the first term containing the temporal dependency of *L* with respect to *W* was truncated and *V ^T^* = *U* from the original recirculation algorithm. Combining the above two equations:

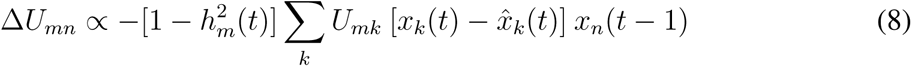

which in the vector notation is:

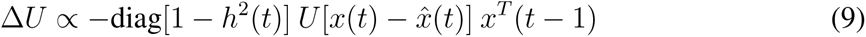

This learning algorithm only involves locally available information: *x*, *x̂*, *h* and *U*. Assuming the input to hidden weights are linear, the term [*h* − *h̃*] in Eq. 4 can be replaced with *U* [*x*(*t*) *− x̂*(*t*)] in Eq. 9, thus making our learning rule for input and output weights approximate the recirculation learning rule described in Eq. 4. As a result, the input and output weights trained from our learning algorithm is approximately the transpose of each other up to scaling (Fig. S7).

#### Recurrent weight

The exact gradient of the recurrent weight (*W*) is:

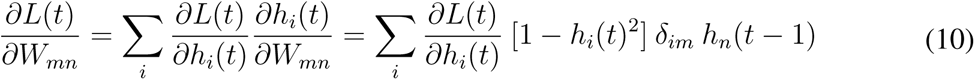

Following the same derivation used to approximate *∂L/∂h* in Eq. 7:

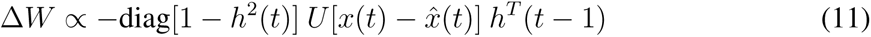

This Hebbian rule between the postsynaptic prediction error and the previous presynaptic input from a recurrent unit is a biologically plausible mechanism for updating the recurrent weights.

The coefficient *σ^′^* = diag(1 *− h*^2^) modulates the learning rate and is only significant around threshold, acting like a gate that restricts weight change to the currently active neurons. In real neurons, this could correspond to backpropagating action potentials that gate synaptic plasticity in dendrites.

## Supplementary figures

**Figure S1:**
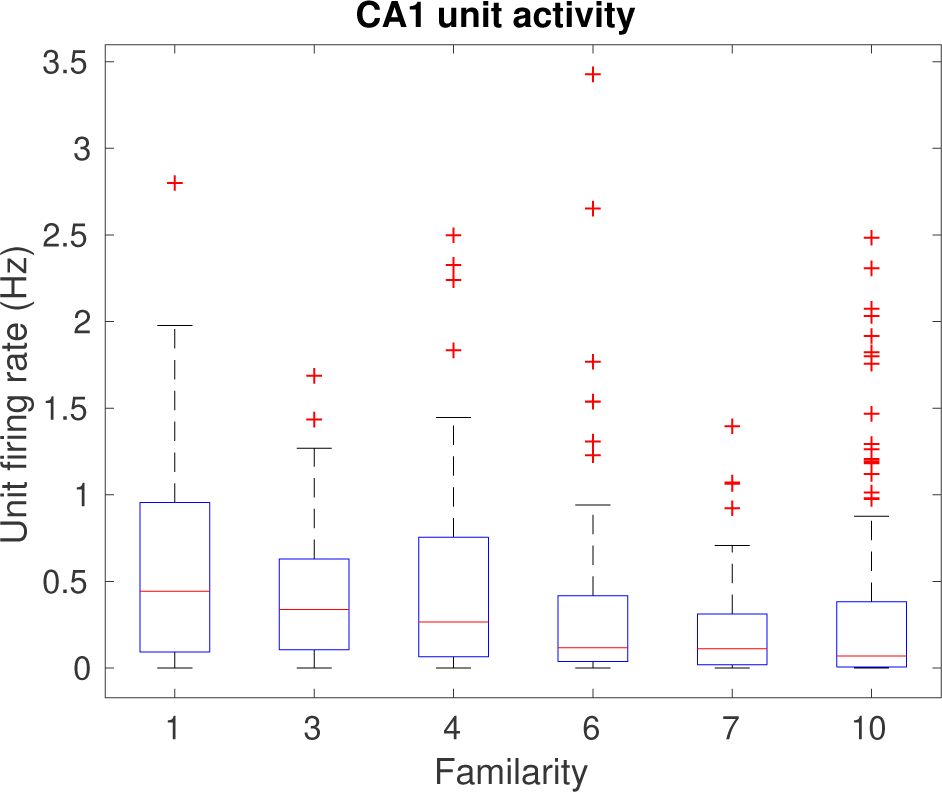
Activities of CA1 neurons decay as the increase of familiarity from CRCNS dataset (Methods).

**Figure S2:**
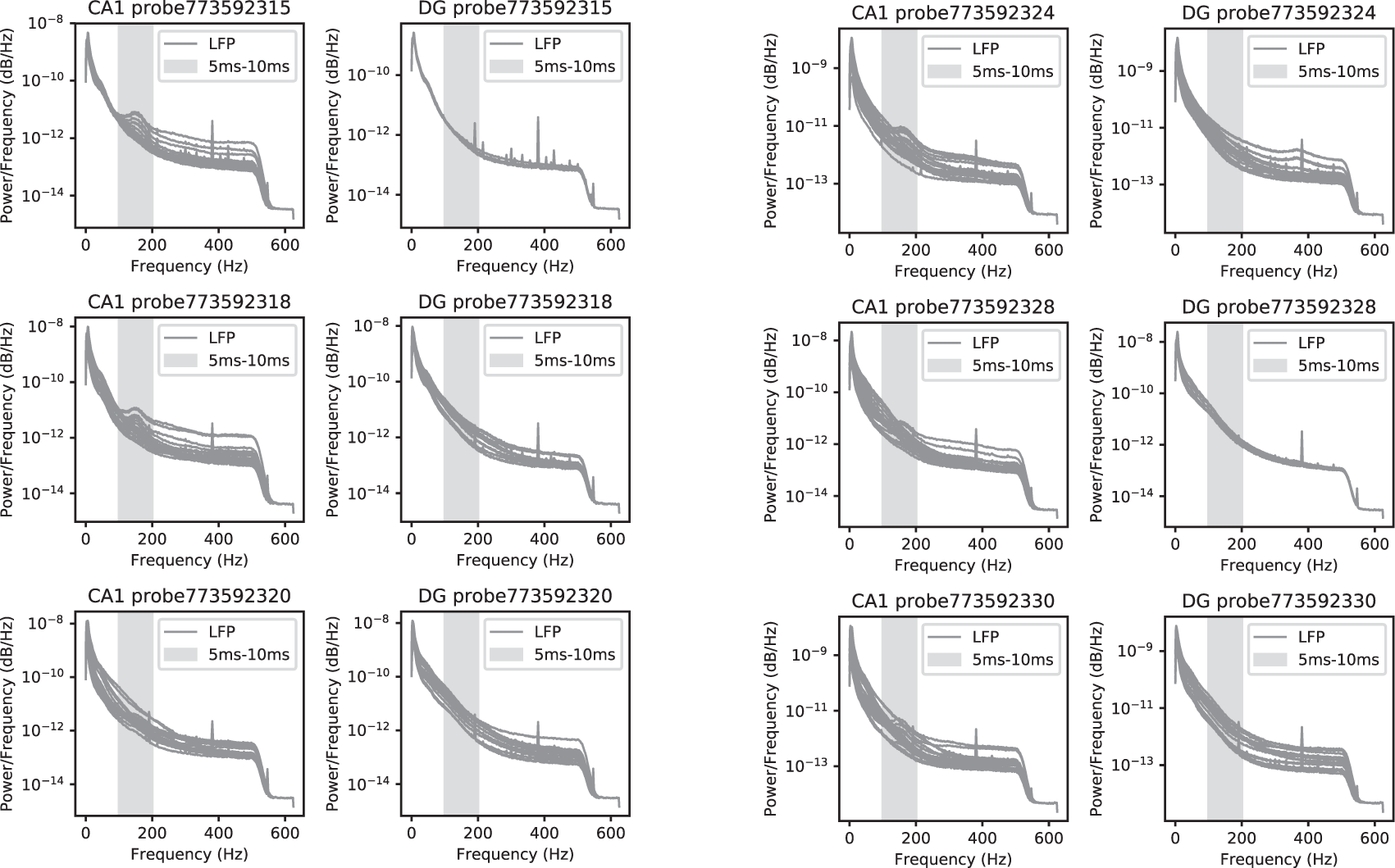
Power Spectrum Density of LFP recorded from probes in CA1 and DG in Session 766640955. Each gray curve stands for one channel on that probe. About one third of CA1 probes showed a power bump from 100Hz to 200 Hz (highlighted by gray background). This indicates that part of CA1 activities oscillate at the period from 5ms to 7ms. This could potentially explain the bimodal distribution in Fig2D.

**Figure S3:**
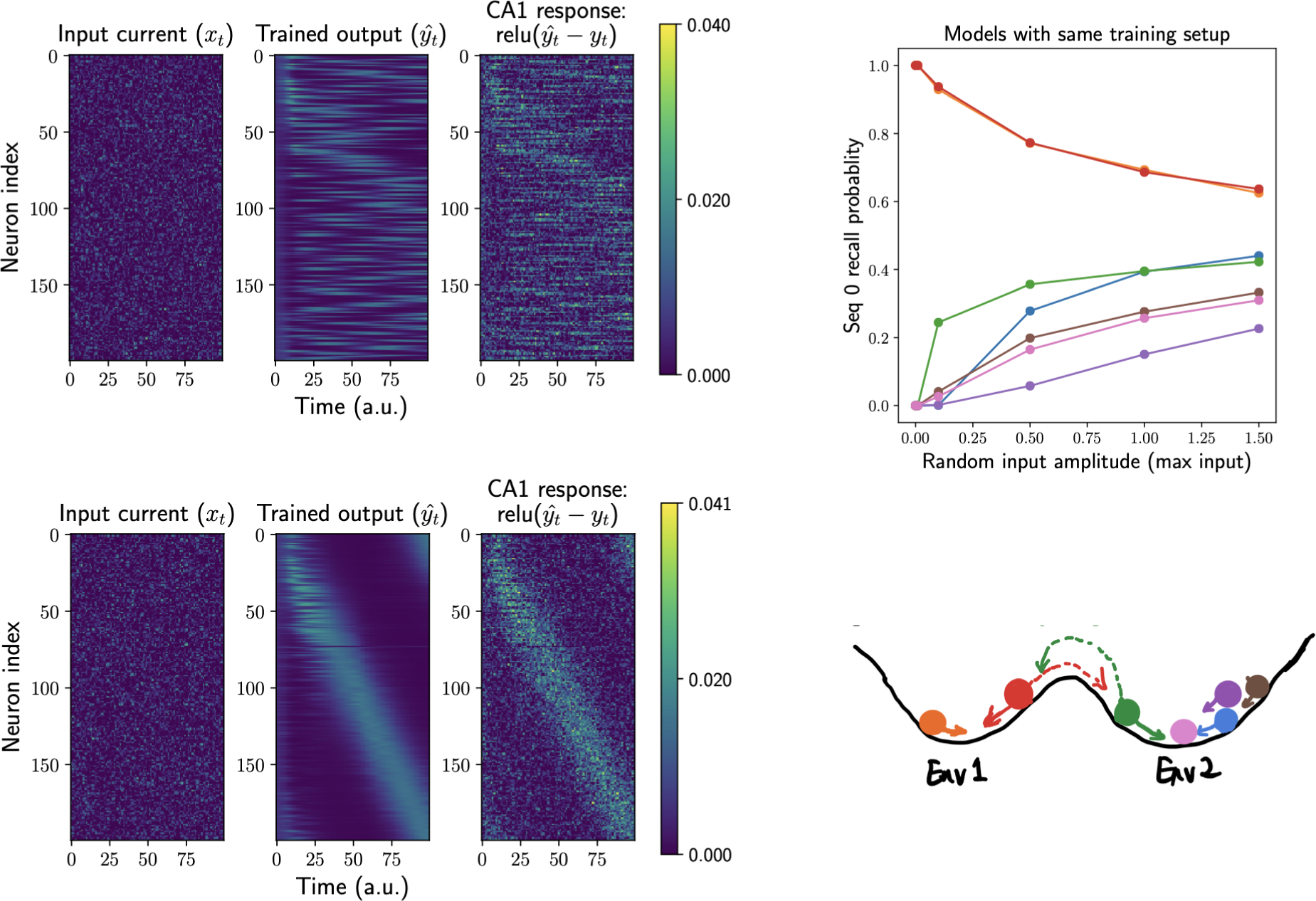
Random recall of one of the remembered sequences. The network was pre- trained to remember two sequences in the same batch. When a random input was given to the network, either one of the sequences would appear as hidden unit activity due to the learnt network structure in Fig.4C (top-left for Env1 reall and bottom-left for Env2 recall). Top-right: The probability of recalling Env1 sequence is either 1 or 0 with different initialization (colored curves) when the input noise level is low. It gradually converges to 0.5 when the input noise level is getting larger. Bottom-right: a schematic showing the energy landscape of two sequences. A larger noise would help push the network to jump between states.

**Figure S4:**
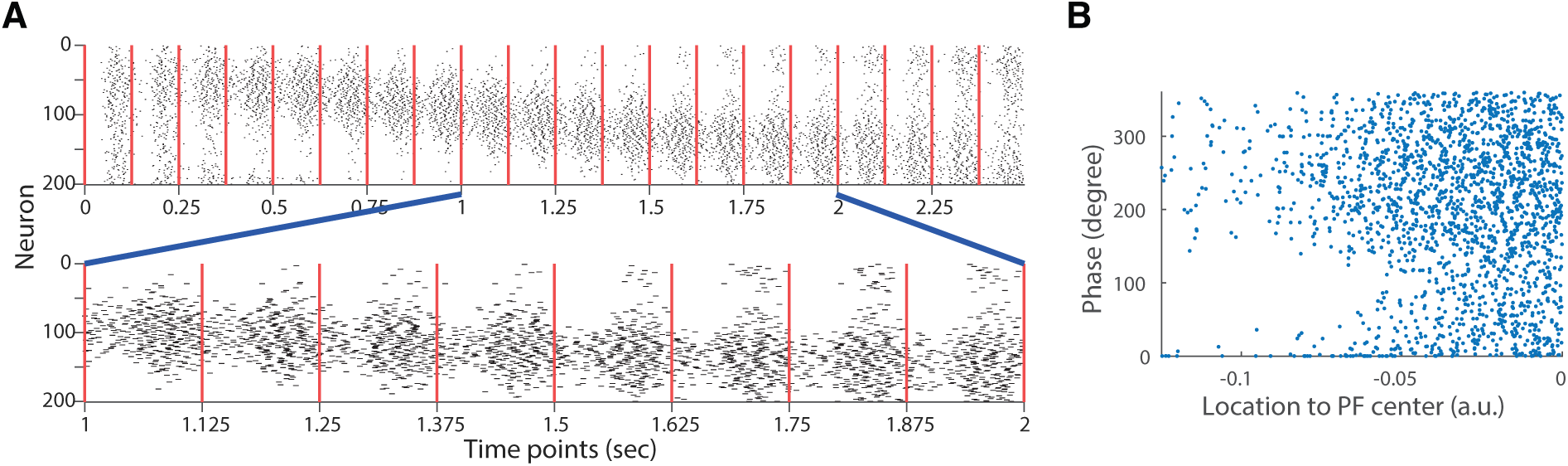
Phase precession of place cell spikes given the structured recurrent weight after training. (A) Spike train recorded from a network of LIF neurons. Red vertical lines mark the trough of 8Hz population activity (i.e. phase=180 degree) (B) Spike phases (relative to 8Hz population activity) of all CA3 neurons plotted against their relative location in a place field.

**Figure S5:**
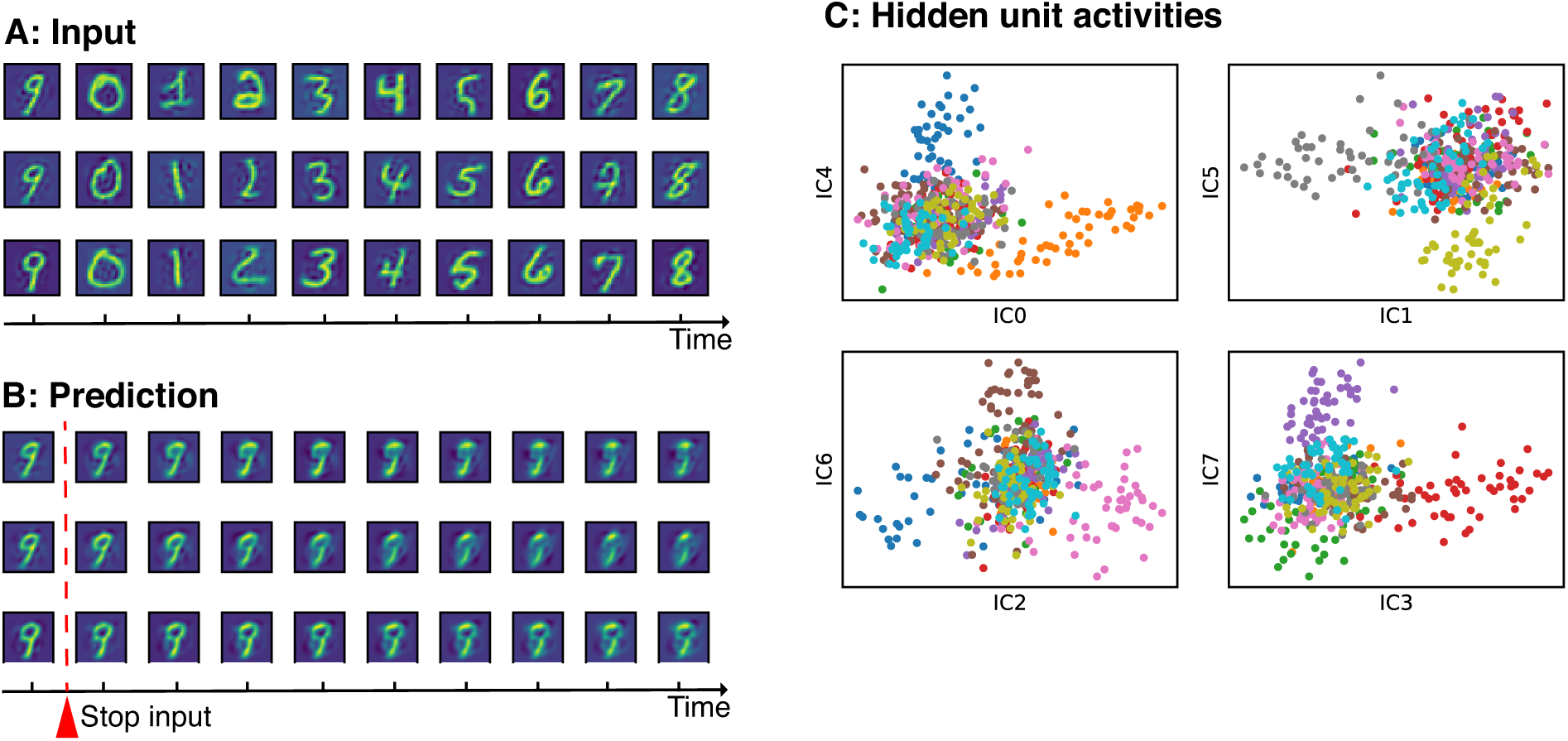
Control using non-predictive loss function. Similar to Fig. 6ABD except that the network was trained using current loss function as a control.

**Figure S6:**
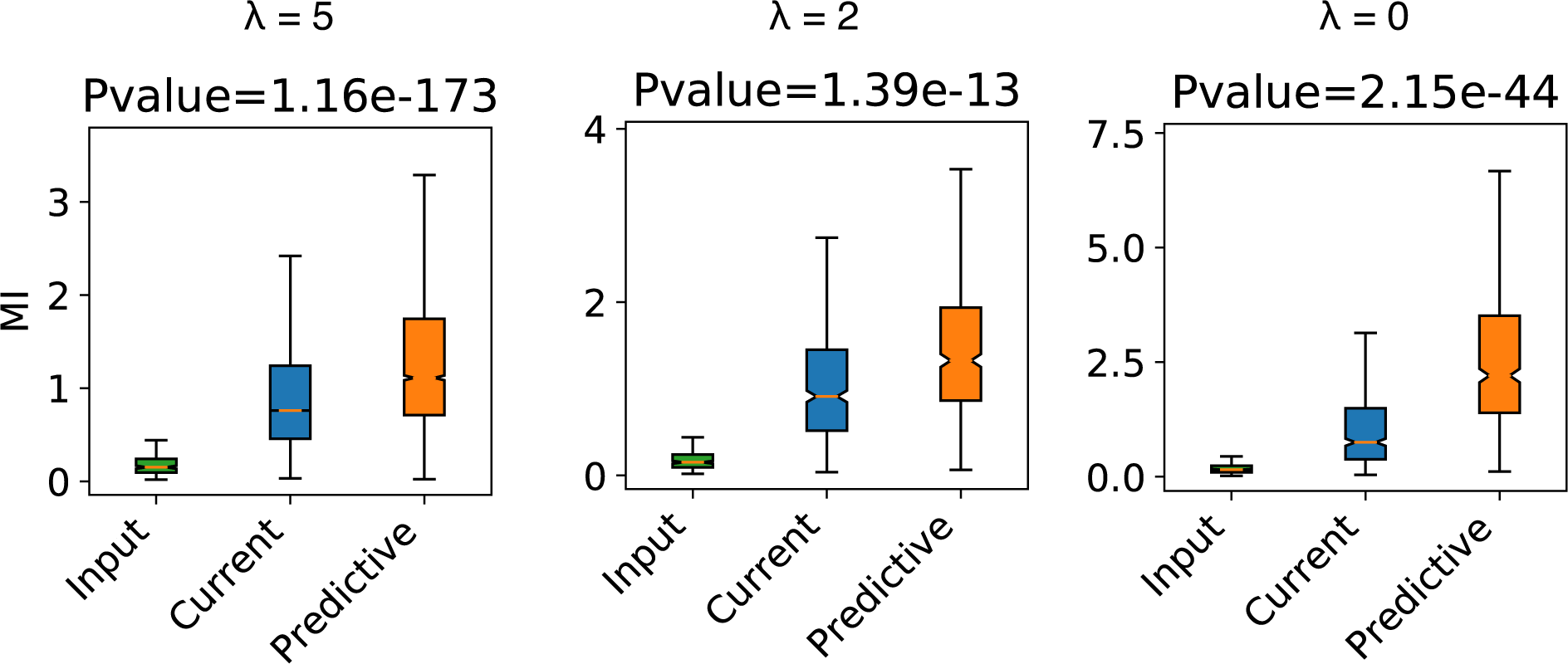
Mutual information of recurrent units trained using different regularization strength (*λ*). P value in the title refers to the comparison (t-test) between units from current network controls and predictive networks. For *λ* = 5, we trained 10 repetitive networks while for the other two, only one representative network was trained.

**Figure S7:**
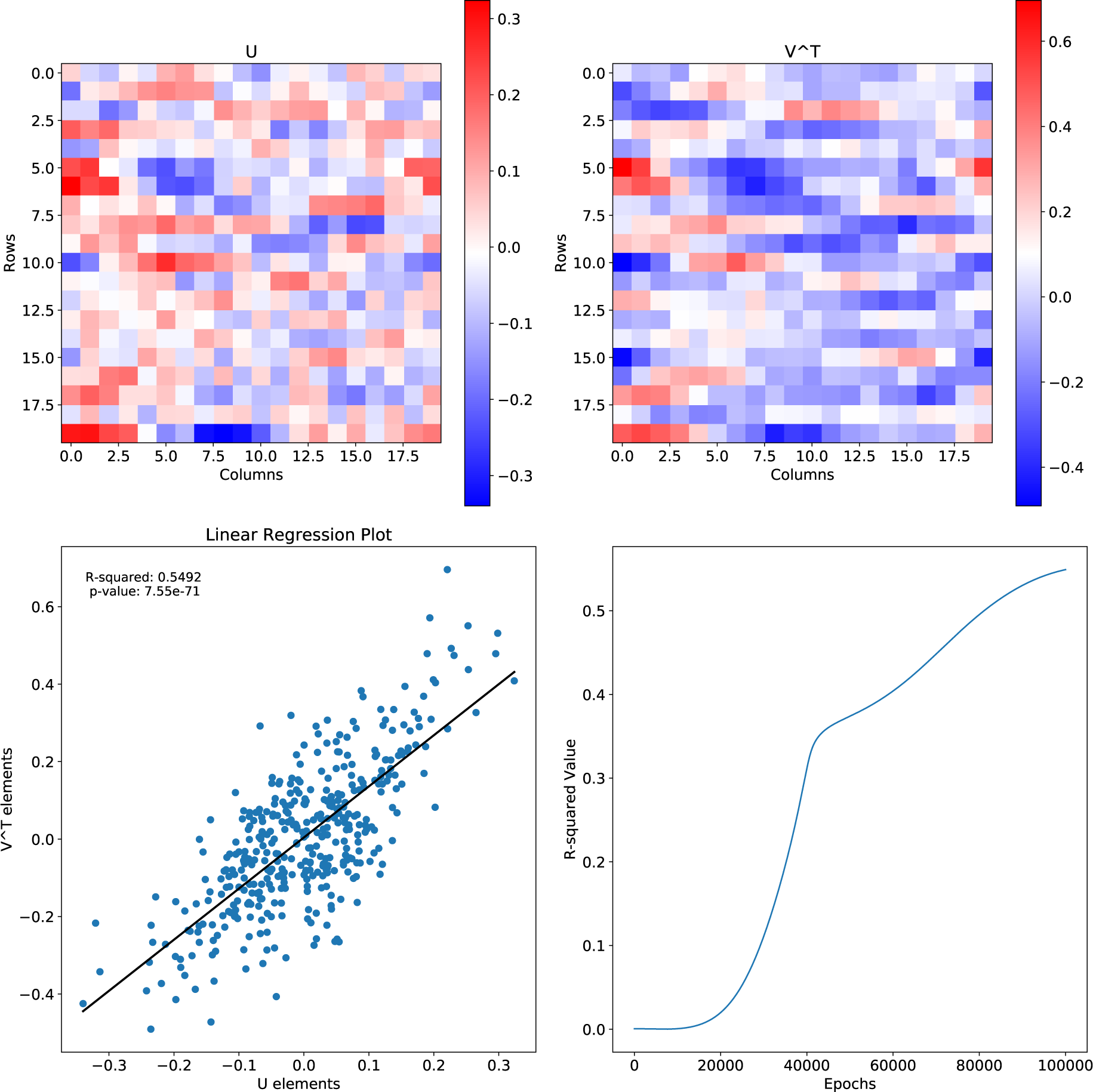
Convergence of input weights *U* (Upper left) and the output weights *V ^T^* (Upper right) after training with predictive recirculation for the simulations in (Fig. 7). Lower left: Scatterplot between entries in *U* and the corresponding entries in *V ^T^*. Linear regression reveals a strong correlation (*r*^2^ = 0.55*, p* = 7.55 *×* 10*^−^*^71^) indicating that these two matrices are approximately the same up to scaling. Lower right: R squared between *U* and *V ^T^* increases with training epochs.

## Footnotes

1 Dan Feldman, personal communication

## Notes

### Competing Interest Statement

The authors have declared no competing interest.

### Summary of Updates

Added Fig.3, 4 and 7. Change a large amount of text

## References

Ackley, D. H., Hinton, G. E., & Sejnowski, T. J. (1985). A learning algorithm for boltzmann machines. Cognitive science, 9, 147–169.

Aronov, D., Nevers, R., & Tank, D. W. (2017). Mapping of a non-spatial dimension by the hippocampal–entorhinal circuit. Nature, 543, 719–722.

Basu, J. & Siegelbaum, S. A. (2015). The corticohippocampal circuit, synaptic plasticity, and memory. Cold Spring Harbor perspectives in biology, 7, a021733.

Benna, M. K. & Fusi, S. (2021). Place cells may simply be memory cells: Memory compression leads to spatial tuning and history dependence. Proceedings of the National Academy of Sciences, 118, e2018422118.

Bi, G.-q. & Poo, M.-m. (1998). Synaptic modifications in cultured hippocampal neurons: dependence on spike timing, synaptic strength, and postsynaptic cell type. Journal of neuroscience, 18, 10464–10472.

Bittner, K. C., Milstein, A. D., Grienberger, C., Romani, S., & Magee, J. C. (2017). Behavioral time scale synaptic plasticity underlies ca1 place fields. Science, 357, 1033–1036.

Bryson, A. E. (1961). A gradient method for optimizing multi-stage allocation processes. Proceedings of the Harvard Univ. Symposium on digital computers and their applications.

Buzsáki, G. & Tingley, D. (2018). Space and time: The hippocampus as a sequence generator. Trends in cognitive sciences, 22, 853–869.

Chandra, S., Sharma, S., Chaudhuri, R., & Fiete, I. (2023). High-capacity flexible hippocampal associative and episodic memory enabled by prestructured”spatial”representations. bioRxiv, pp. 2023–11.

Davoudi, H. & Foster, D. J. (2019). Acute silencing of hippocampal ca3 reveals a dominant role in place field responses. Nature neuroscience, 22, 337–342.

Dong, C., Madar, A. D., & Sheffield, M. E. (2021). Distinct place cell dynamics in ca1 and ca3 encode experience in new environments. Nature communications, 12, 2977.

Dragoi, G. & Tonegawa, S. (2011). Preplay of future place cell sequences by hippocampal cellular assemblies. Nature, 469, 397–401.

Duncan, K., Ketz, N., Inati, S. J., & Davachi, L. (2012). Evidence for area ca1 as a match/mismatch detector: A high-resolution fmri study of the human hippocampus. Hippocampus, 22, 389–398.

Eichenbaum, H., Kuperstein, M., Fagan, A., & Nagode, J. (1987). Cue-sampling and goalapproach correlates of hippocampal unit activity in rats performing an odor-discrimination task. Journal of Neuroscience, 7, 716–732.

Eliasmith, C. (2007). Attractor network. Scholarpedia, 2, 1380. revision #91016.

Fang, C., Aronov, D., Abbott, L., & Mackevicius, E. L. (2023). Neural learning rules for generating flexible predictions and computing the successor representation. Elife, 12, e80680.

Gardner, R. J., Hermansen, E., Pachitariu, M., Burak, Y., Baas, N. A., Dunn, B. A., Moser, M.-B., & Moser, E. I. (2022). Toroidal topology of population activity in grid cells. Nature, 602, 123–128.

Gauthier, J. L. & Tank, D. W. (2018). A dedicated population for reward coding in the hippocampus. Neuron, 99, 179–193.

George, D., Rikhye, R. V., Gothoskar, N., Guntupalli, J. S., Dedieu, A., & Lázaro-Gredilla, M. (2021). Clone-structured graph representations enable flexible learning and vicarious evaluation of cognitive maps. Nature communications, 12, 2392.

Gökçe, O., Bonhoeffer, T., & Scheuss, V. (2016). Clusters of synaptic inputs on dendrites of layer 5 pyramidal cells in mouse visual cortex. Elife, 5, e09222.

Grienberger, C. & Magee, J. C. (2022). Entorhinal cortex directs learning-related changes in ca1 representations. Nature, 611, 554–562.

Haeusler, S. & Wolfgang, M. (2007). A statistical analysis of information-processing properties of lamina-specific cortical microcircuit models. Cerebral Cortex, 17, 149–162.

Hinton, G. E. & McClelland, J. (1987). Learning representations by recirculation. In Neural information processing systems.

Hinton, G. E., Sejnowski, T. J., et al. (1986). Learning and relearning in boltzmann machines. Parallel distributed processing: Explorations in the microstructure of cognition, 1, 2.

Hopfield, J. J. (1982). Neural networks and physical systems with emergent collective computational abilities. Proceedings of the national academy of sciences, 79, 2554–2558.

Isomura, T. & Toyoizumi, T. (2021). Dimensionality reduction to maximize prediction generalization capability. Nature Machine Intelligence, 3, 434–446.

Jaeger, H. & Haas, H. (2004). Harnessing nonlinearity: Predicting chaotic systems and saving energy in wireless communication. science, 304, 78–80.

Keller, T. A., Muller, L., Sejnowski, T., & Welling, M. (2023). Traveling waves encode the recent past and enhance sequence learning. arXiv preprint arXiv:2309.08045.

Khona, M. & Fiete, I. R. (2022). Attractor and integrator networks in the brain. Nature Reviews Neuroscience, 23, 1–23.

Kim, R., Li, Y., & Sejnowski, T. J. (2019). Simple framework for constructing functional spiking recurrent neural networks. Proceedings of the national academy of sciences, 116, 22811–22820.

Kingma, D. P. & Welling, M. (2019). An introduction to variational autoencoders. arXiv preprint arXiv:1906.02691.

Klausberger, T. (2009). Gabaergic interneurons targeting dendrites of pyramidal cells in the ca1 area of the hippocampus. European Journal of Neuroscience, 30, 947–957.

Knight, R. T. (1996). Contribution of human hippocampal region to novelty detection. Nature, 383, 256–259.

Kumaran, D. & Maguire, E. A. (2006). An unexpected sequence of events: mismatch detection in the human hippocampus. PLoS biology, 4.

Lee, I., Rao, G., & Knierim, J. J. (2004a). A double dissociation between hippocampal subfields: differential time course of ca3 and ca1 place cells for processing changed environments. Neuron, 42, 803–815.

Lee, I., Yoganarasimha, D., Rao, G., & Knierim, J. J. (2004b). Comparison of population coherence of place cells in hippocampal subfields ca1 and ca3. Nature, 430, 456–459.

Leung, L. S., Roth, L., & Canning, K. J. (1995). Entorhinal inputs to hippocampal ca1 and dentate gyrus in the rat: a current-source-density study. Journal of Neurophysiology, 73, 2392–2403.

Leutgeb, S., Leutgeb, J. K., Treves, A., Moser, M.-B., & Moser, E. I. (2004). Distinct ensemble codes in hippocampal areas ca3 and ca1. Science, 305, 1295–1298.

Lisman, J. E. & Grace, A. A. (2005). The hippocampal-vta loop: controlling the entry of information into long-term memory. Neuron, 46, 703–713.

Lotter, W., Kreiman, G., & Cox, D. (2016). Deep predictive coding networks for video prediction and unsupervised learning. arXiv preprint arXiv:1605.08104.

Lotter, W., Kreiman, G., & Cox, D. (2020). A neural network trained for prediction mimics diverse features of biological neurons and perception. Nature Machine Intelligence, 2, 210–219.

Markov, N. T., Ercsey-Ravasz, M., Van Essen, D. C., Knoblauch, K., Toroczkai, Z., & Kennedy, H. (2013). Cortical high-density counterstream architectures. Science, 342, 1238406.

Markram, H., Lübke, J., Frotscher, M., & Sakmann, B. (1997). Regulation of synaptic efficacy by coincidence of postsynaptic aps and epsps. Science, 275, 213–215.

McNamee, D. C., Stachenfeld, K. L., Botvinick, M. M., & Gershman, S. J. (2021). Flexible modulation of sequence generation in the entorhinal–hippocampal system. Nature Neuroscience, 24, 851–862.

Mizuseki, K., Diba, K., Pastalkova, E., Teeters, J., Sirota, A., & Buzsáki, G. (2014). Neurosharing: large-scale data sets (spike, lfp) recorded from the hippocampal-entorhinal system in behaving rats. F1000Research, 3, 98.

Mizuseki, K., Sirota, A., Pastalkova, E., Diba, K., & Buzsáki, G. (2013). Multiple single unit recordings from different rat hippocampal and entorhinal regions while the animals were performing multiple behavioral tasks. CRCNS org.

Nieh, E. H., Schottdorf, M., Freeman, N. W., Low, R. J., Lewallen, S., Koay, S. A., Pinto, L., Gauthier, J. L., Brody, C. D., & Tank, D. W. (2021). Geometry of abstract learned knowledge in the hippocampus. Nature, 595, 80–84.

O’Keefe, J. & Nadel, L. (1978). The hippocampus as a cognitive map. (Oxford university press).

O’Keefe, J. & Recce, M. L. (1993). Phase relationship between hippocampal place units and the eeg theta rhythm. Hippocampus, 3, 317–330.

Ólafsdóttir, H. F., Bush, D., & Barry, C. (2018). The role of hippocampal replay in memory and planning. Current Biology, 28, R37–R50.

Pastalkova, E., Itskov, V., Amarasingham, A., & Buzsaki, G. (2008). Internally generated cell assembly sequences in the rat hippocampus. Science, 321, 1322–1327.

Patel, J., Fujisawa, S., Berényi, A., Royer, S., & Buzsáki, G. (2012). Traveling theta waves along the entire septotemporal axis of the hippocampus. Neuron, 75, 410–417.

Priestley, J. B., Bowler, J. C., Rolotti, S. V., Fusi, S., & Losonczy, A. (2022). Signatures of rapid plasticity in hippocampal ca1 representations during novel experiences. Neuron.

Rao, R. P. & Ballard, D. H. (1999). Predictive coding in the visual cortex: a functional interpretation of some extra-classical receptive-field effects. Nature neuroscience, 2, 79–87.

Recanatesi, S., Farrell, M., Lajoie, G., Deneve, S., Rigotti, M., & Shea-Brown, E. (2021). Predictive learning as a network mechanism for extracting low-dimensional latent space representations. Nature communications, 12, 1–13.

Rumelhart, D. E., Hinton, G. E., & Williams, R. J. (1986). Learning representations by backpropagating errors. nature, 323, 533–536.

Sabatini, B. & Regehr, W. (1999). Timing of synaptic transmission. Annual review of physiology, 61, 521–542.

Schapiro, A. C., Turk-Browne, N. B., Botvinick, M. M., & Norman, K. A. (2017). Complementary learning systems within the hippocampus: a neural network modelling approach to reconciling episodic memory with statistical learning. Philosophical Transactions of the Royal Society B: Biological Sciences, 372, 20160049.

Sejnowski, T. J. (2023). Large language models and the reverse turing test. Neural computation, 35, 309–342.

Shin, J., Lee, H.-W., Jin, S.-W., & Lee, I. (2022). Subtle visual change in a virtual environment induces heterogeneous remapping systematically in ca1, but not ca3. Cell Reports, 41, 111823.

Siegle, J. H., Jia, X., Durand, S., Gale, S., Bennett, C., Graddis, N., Heller, G., Ramirez, T. K., Choi, H., Luviano, J. A., et al. (2021). Survey of spiking in the mouse visual system reveals functional hierarchy. Nature, 592, 86–92.

Skaggs, W., Mcnaughton, B., & Gothard, K. (1992). An information-theoretic approach to deciphering the hippocampal code. Advances in neural information processing systems, 5.

Stachenfeld, K. L., Botvinick, M. M., & Gershman, S. J. (2017). The hippocampus as a predictive map. Nature neuroscience, 20, 1643–1653.

Tsodyks, M. V., Skaggs, W. E., Sejnowski, T. J., & McNaughton, B. L. (1996). Population dynamics and theta rhythm phase precession of hippocampal place cell firing: a spiking neuron model. Hippocampus, 6, 271–280.

Vaswani, A., Shazeer, N., Parmar, N., Uszkoreit, J., Jones, L., Gomez, A. N., Kaiser, L. u., & Polosukhin, I. (2017). Attention is all you need. In Advances in Neural Information Processing Systems, I. Guyon, U. V. Luxburg, S. Bengio, H. Wallach, R. Fergus, S. Vishwanathan, & R. Garnett, eds., vol. 30. (Curran Associates, Inc.).

Watabe-Uchida, M., Eshel, N., & Uchida, N. (2017). Neural circuitry of reward prediction error. Annual Review of Neuroscience, 40, 373—394.

Werbos, P. J. (1990). Backpropagation through time: what it does and how to do it. Proceedings of the IEEE, 78, 1550–1560.

Whittington, J. C., Muller, T. H., Mark, S., Chen, G., Barry, C., Burgess, N., & Behrens, T. E. (2020). The tolman-eichenbaum machine: Unifying space and relational memory through generalization in the hippocampal formation. Cell, 183, 1249–1263.

Wikipedia (2021). Recursive bayesian estimation — Wikipedia, the free encyclopedia. [Online; accessed 8-May-2022].

Wilson, M. A. & McNaughton, B. L. (1994). Reactivation of hippocampal ensemble memories during sleep. Science, 265, 676–679.

Zhang, K. (1996). Representation of spatial orientation by the intrinsic dynamics of the head-direction cell ensemble: a theory. Journal of Neuroscience, 16, 2112–2126.

